# The regulatory genome of the malaria vector *Anopheles gambiae*: integrating chromatin accessibility and gene expression

**DOI:** 10.1101/2020.06.22.164228

**Authors:** José L. Ruiz, Lisa C. Ranford-Cartwright, Elena Gómez-Díaz

**Affiliations:** Instituto de Parasitología y Biomedicina López-Neyra (IPBLN), Consejo Superior de Investigaciones Científicas, 18016, Granada, Spain; Institute of Biodiversity, Animal Health and Comparative Medicine, College of Medical, Veterinary and Life Science, University of Glasgow, Glasgow G12 8QQ, UK

## Abstract

*Anopheles gambiae* mosquitoes are primary human malaria vectors, but we know very little about mechanisms of transcriptional regulation. We profiled chromatin accessibility by ATAC-seq in laboratory-reared *An. gambiae* mosquitoes experimentally infected with the human malaria parasite *Plasmodium falciparum*. By integrating ATAC-seq, RNA-seq and ChIP-seq data we showed a positive correlation between accessibility at promoters and introns, gene expression and active histone marks. By comparing expression and chromatin structure patterns in different tissues, we were able to infer cis-regulatory elements controlling tissue specific gene expression and to predict the in vivo binding sites of relevant transcription factors. The ATAC-seq assay also allowed the precise mapping of active regulatory regions, including novel transcription start sites and enhancers that annotate to mosquito immune-response genes. This study is important not only for advancing our understanding of mechanisms of transcriptional regulation in the mosquito vector of human malaria, but the information is of great potential for developing new mosquito-control and anti-malaria strategies.

## INTRODUCTION

Chromatin structure is the basal element determining dynamic regulatory landscapes, i.e. the set of regulatory sequences, and the proteins binding to them, that control the definition of phenotypes during development, and in response to the external environment, in metazoa (1). Cis-regulatory elements are regions of non-coding DNAs capable of regulating transcription. For example, accessibility at promoters allows for the interaction of transcription factors (TFs) with their cognate motifs, recruiting other co-factors involved in chromatin remodeling and transcriptional activation, that enables the spatiotemporal control of gene expression (2-4). Additionally, transcriptional enhancers work in concert with the core promoter in regulating gene expression, acting as a scaffold for the recruitment of transcription factors and chromatin-modifying enzymes (5-8). Other relevant regulatory regions are insulators, that typically work in long-range distances and contain binding sites for specific TFs, such as CTCF (9). Chromatin structure and accessibility at these regulatory regions can also influence alternative splicing (10). The regulation of this process often involves intronic or exonic cis-regulatory elements that are bound by DNA-binding proteins, that interfere with RNA polymerase II transcriptional elongation or associate with enhancers or silencers in a time- and tissue-specific manner (11-14). Many splicing regulatory elements have been identified in mammals (15-17), only a few have been described in *Drosophila* (18,19), and, in general, studies examining the relationship between the dynamics of chromatin accessibility at cis-regulatory sequences mediating alternative splicing events are scarce in invertebrates (20-22).

Based on these fundamental principles, variable levels of chromatin accessibility at regulatory regions are expected to reflect the level of transcriptional activity at a given tissue or condition and time-point. It is also expected that these regulatory sites display a typical pattern of histone post-translational modifications (hPTMs) characteristic of active chromatin (23). As a consequence, chromatin accessibility can be used as a proxy to globally identify active promoters and enhancers, and to predict gene activity (24,25). By profiling genome-wide chromatin accessibility in several model organisms, such as the fruit fly *Drosophila melanogaster*, recent studies have mapped thousands of cis- and trans-regulatory elements, defining their functional roles in the regulation of genes involved in processes such as development, physiology and disease (26,27). For example, there are more than 40,000 *Drosophila* enhancers and target genes described in the EnhancerAtlas database (27). These enhancers, in general, have been shown to modulate the transcription levels of several target genes, regardless of orientation or distance to the target promoters (5,28,29), and they can be located between and within genes, i.e. within introns or exons (6,26,30).

Compared to the knowledge accumulated on transcriptional regulation in the fruit fly, little is known about the regulatory genome of other insects, such as mosquitoes. This is despite the major role of these arthropod vectors in the transmission of important human infectious diseases. Amongst mosquito-borne diseases, malaria is the deadliest and the one with the highest global health and economic burden (31). Human malaria is transmitted by *Anopheles* mosquitoes, with members of the species complex *An. gambiae* recognised as the main vectors in Africa (32). Controlling and targeting vector populations is key in the ongoing efforts to fight malaria, but further progress towards alternative molecular-based approaches has been hampered by the lack of epigenetic and functional genomic studies in mosquitoes (33,34). Indeed, the regulatory genome of *Anopheles* remains practically unexplored and the regulatory networks of most genes in the *An. gambiae* genome are unknown, including those genes involved in important functions such as mosquito insecticide resistance and immunity. The vast majority of cis-regulatory sequences reported to date in mosquitoes are computational predictions without experimental verification (35-42). For example, in relation to promoters and TSSs in *Anopheles*, most 5’ UTRs have been mapped already. However, in *An. gambiae* a large proportion of genes (2,890 out of 13,057, or 22.1%, considering updated genome annotations) still do not have 5’ UTRs annotated, a proportion that is in the same range to that reported in *D. melanogaster* (43). More strikingly, less than a dozen enhancer sequences have been experimentally validated in *An. gambiae* (38,44), a negligible number compared with the functionally annotated enhancers that are publicly available in several *Drosophila* data bases (26,27,45,46). Other aspects of the genomics of mosquitoes that remains little studied from a mechanistic perspective, and that might be related to the function of unknown cis-regulatory elements, are isoform switching and chromatin-associated mechanisms of regulation.

To fill this important gap, here we used the Assay of Transposase-Accessible Chromatin by sequencing (ATAC-seq) (47,48) to characterize the *An. gambiae* regulatory landscape. A comprehensive and integrative examination of genome-wide chromatin accessibility patterns was necessary to further characterize mosquito gene regulatory networks *in vivo*. This analysis of chromatin accessibility allows for the genome-wide identification of promoter regions and enhancers, as well as the prediction TF binding events (2). In the research presented here, we performed ATAC-seq and RNA-seq analyses of both *An. gambiae* midguts and salivary glands infected with *P. falciparum*, and integrated such datasets with ChIP-seq data for various histone modifications (H3K9ac, H3K4me3, H3K27ac and H3K9m3) (42). Our aim was to provide a more complete annotation of the regulatory genome of a major mosquito vector of human malaria, and to add new insights into mechanisms controlling functional gene expression. We report a genome-wide association between chromatin accessibility, epigenetic states and tissue-specific regulation of transcription. Analyses of DNA-binding motif enrichment of active regulatory regions allowed us to predict binding sites similar to several *Drosophila* TFs, many of which have been functionally validated. Furthermore, we provide a comprehensive map of enhancer-, TSS- and CTCF-like novel regulatory sequences, which are conserved with those previously characterized in the fruit fly, and appear to be active in the mosquito.

## MATERIAL AND METHODS

### Mosquito rearing and experimental infections

Five-day old female *An. gambiae* s.s. Kisumu mosquitoes from a genetically outbred laboratory colony in the University of Glasgow were used for experimental infections. Mosquitoes were maintained under standard insectary conditions (27□±□2 °C, 80□±□5% relative humidity, 12:12 LD). Females were fed through membrane feeders on blood containing gametocytes of *P. falciparum* clone 3D7, prepared according to standard protocols (49). Thereafter the mosquitoes were given access to a solution of 5% glucose/ 0.05% 4-aminobenzoic acid *ad libitum*. Three independent experimental infections (Infections 1, 2 and 3) were carried out. Prevalence (percentage of infected mosquitoes) and intensity of infection (median number of oocysts) are described in Supplementary Table S1. We performed dissection of midguts (MGs) on adults at 7 days post-infection and of salivary glands (SGs) at 14 days post-blood meal.

### ATAC-seq library preparation and sequencing

We performed the ATAC-seq protocol using fresh MGs and SGs from ∼20 individual mosquitoes from two independent infections (Supplementary Table S1). Mosquito tissues were resuspended in lysis buffer (47,48) to permeabilize membranes. The nuclei pellet was resuspended in the transposition reaction mix (25 μl of 2×TD Buffer, 1.25 μl of Tn5 Transposase and 23.75 μl of nuclease free water), and incubated for 30 min at 37°C. All samples were purified using the Qiagen MiniElute Kit. Library amplification was carried out with 2X KAPA HiFi mix and 1.25 μM of Nextera primers (47,48). The optimal cycle number was determined by qPCR with conditions as originally described in (47,48). ATAC-seq libraries were sequenced at BGI (China) with an Illumina HiSeq4000 sequencer to obtain 25-37 M of 2×50 bp paired-end reads (Supplementary Table S2).

### RNA isolation, RNA-seq library preparation and sequencing

We prepared RNA-seq libraries from *P. falciparum-*infected midguts and salivary glands obtained in two independent experimental infections (2 and 3; Supplementary Table S1). Immediately after dissection, tissues were stored in TRIzol (Invitrogen) and frozen at −80°C until subsequent processing. Total RNA was extracted from a pool of ∼□30 midguts and a pool of ∼60 salivary glands from 30 mosquitoes using the TRIzol manufacturer protocol. RNA concentration was quantified using a Qubit® 2.0 Fluorometer, and RNA integrity was determined with an Agilent 2100 Bioanalyzer. We used the Ovation® Universal RNA-seq System (Nugen Technologies) for strand specific RNA-seq library construction following the manufacturer’s instructions. Custom primers specific to mosquito ribosomal sequences were designed to reduce the percentage of ribosomal reads in the sample and the ribo-depletion step was incorporated into the standard workflow (Supplementary Table S3). Libraries were sequenced at Cabimer (Spain) using an Illumina NextSeq500 for both 2×150 bp paired-end and 1×75 bp single-end reads.

### ATAC-seq data processing and analyses

We conducted ATAC-seq data analyses according to the recommendations of the ENCODE Pipeline (https://www.encodeproject.org/atac-seq/, (50)). First, raw reads were trimmed 20 bases from the 3′ end of each read (−3 20), and aligned to the *An. gambiae* PEST genome (AgamP4) with Bowtie2 (51) (v2.4.1) using default parameters, except for –no-unal –no-mixed -X 2000. We then applied a MAPQ score threshold (10) and sorted and deduplicated the reads using SAMtools (52) (v1.10). To adjust the known bias and ensure the mapping of Tn5 cutting sites, we shifted aligned reads +4 bp for + strands and −5 bp for – strands with ATACSeqQC (53) (v1.10). We removed not properly-paired reads and extracted nucleosome-free fragments with a size threshold of 130 bp (SAMtools). We performed peak-calling on nucleosome-free reads with MACS2 (54) (v.2.1.2) *callpeak* module and the following parameters: -f BAMPE -g 273109044 -q 0.01 -B --keep-dup all -- nomodel --nolambda. We refer to ATAC-seq peaks as Tn5 hypersensitive sites (THSs). To identify THSs unique or common across samples, we used BEDTools *intersect* (55) requiring a minimum overlap of 51% (-f/-F 0.51). We annotated THSs to genomic features combining HOMER (56) (v4.11) and ChIPseeker (57) (v1.22) (see Supplementary Methods). We used the AgamP4.12 gene set (58) and considered the first coordinate of the 5’ UTRs as the TSSs. For genes without an annotated 5’ UTR we took the translation start site (ATG codon) as the reference point. In either case, the 1 Kb upstream window was considered as the putative promoter region. The mean length of the annotated 5’ UTRs was 253 bp, and the higher density in the distribution of values is ∼100 bp, so even if some genes do not have annotated 5’ UTRs, we expect a 1 Kb window to capture the whole promoter (Supplementary Figure S5A). Metadata for the annotated genes was obtained from the Gene Ontology (GO), Kyoto Encyclopedia of Genes and Genomes (KEGG) and Pfam databases using DAVID (59) and PANTHER (60), and the WikiPathways (61) and ImmunoDB (62) databases (Supplementary Table S4). To quantify the ATAC-seq nucleosome-free signal enrichment at genomic regions of interest, we used BEDTools *intersect* -c. The read counts were normalized (RPKM) and we added a pseudocount (0.1) when required to get finite values. We categorized genes into high, medium or low groups based on promoter chromatin accessibility with threshold values determined by dividing the signal in three quantile groups according to their means (*Hmisc::cut2* R function).

We extracted mononucleosomal reads applying a 171-256 bp size threshold (SAMtools) and predicted nucleosome dyads using NucleoATAC (63) (v0.3.4) (see Supplementary Methods).

### RNA-seq data processing and analyses

We trimmed adapters from the raw reads using BBDuk (v38.79) with -tbo -tpe -minlength=35 parameters and removed rRNA contamination using SortMeRNA (64) (v4.2) with default parameters. Apart from the default rRNA databases, we used additional large and small subunit *An. gambiae* and *P. falciparum* rRNA sequences from the SILVA database (65,66) (LSU r132/SSU r138). Cleaned directional RNA-seq reads were mapped against the AgamP4 v2.00 reference genome using STAR (67) (v2.7.3a). Two different sets of reads were available, 2×150 bp and 1×75bp, which were processed in parallel until we combined the raw counts. Raw counts at the gene level were obtained using CoCo (68) (v.0.2.2) and then provided to DESeq2 (69) (v1.26) for normalization (see Supplementary Methods). Correlation and PCA plots by deepTools2 (70) (v3.4.1) showed higher similarity between infections than between tissues and no clustering based on the sequencing approach (Supplementary Figures S5B and C), which is consistent with results using summed counts. Normalized counts were comparable between infections and higher in SGs (Supplementary Figure S5D). We also observed high correlation (∼75%) between the RNA-seq data in this study and data from our previous study (42) (Supplementary Figure S5E). We categorized normalized counts in high, medium or low groups as described above (*Hmisc::cut2* R function). Normalized RNA-seq counts (DESeq2) for each gene are included in Supplementary Table S5.

### Integration of ATAC-seq, RNA-seq and ChIP-seq data

We used ChromHMM (71) (v1.2) to compute genome-wide chromatin state predictions on the *An. gambiae* genome based on ATAC-seq nucleosome-free signal enrichment levels and hPTMs data from our previous study (42) (see Supplementary Methods). This analysis was performed only on midguts for which hPTMs data was available.

For the correlation and integrative analyses of ATAC-seq, ChiP-seq and RNA-seq data, we restricted the analysis to a subset of 8,245 genes that harbour high confidence regulatory regions. We first discarded genes with very close adjacent genes encoded on opposing strands (with upstream promoters likely overlapping) and genes with the gene bodies or putative promoters overlapping or embedded into many other gene bodies or promoters (> 2). Following their categorization into high, medium, low accessibility groups (see above) we identified genes displaying different patterns of regulation: activating when the change in accessibility is in the same direction, and potential repressor events when the promoter is accessible but the gene is weakly expressed or silent.

### Differential ATAC-seq and RNA-seq analyses

We used the DiffBind R package (72) (v2.14) to assess differential chromatin accessibility at given locations between mosquito MGs and SGs. As input for DiffBind, we used the ATAC-seq nucleosome-free reads and the THSs. Infection 1 and Infection 2 were used as biological replicates. Differential gene expression analyses between mosquito MGs and SGs were conducted using the DESeq2, edgeR (73) (v3.28.1) and DREAMSeq (74) (v1.0.4) R packages to identify Differentially Expressed Genes (DEGs) (Supplementary Table S6). We used the IsoformSwitchAnalyzeR R package (75) (v1.8) to analyze differential gene isoforms expression between MGs and SGs. See Supplementary Methods for more information and details on these analyses.

### Characterization of novel regulatory elements

To map active regulatory sites, we used *An. gambiae* enhancers predicted computationally by others from *Drosophila* enhancers (N=1,628), or from *Drosophila* enhancer motifs (N=51) (37,38), as well a few *Anopheles* enhancers identified previously by STARR-seq (N=6) (44) (see Supplementary Methods). Next, to generate a set of novel candidates, we downloaded *D. melanogaster* collections of enhancers (27), including some activity-based enhancer-target gene assignments (26). We then used the UCSC LiftOver tool webserver (https://genome.ucsc.edu/cgi-bin/hgLiftOver, (76)), which uses homology data between species, to transfer coordinates to *An. gambiae* (see Supplementary Methods). In both sets of enhancers, we checked the overlap of the novel enhancer-like elements with our THSs using BEDTools *intersect* and incorporated the annotation of the THSs (see above and Supplementary Methods). The target genes for the previous enhancers were obtained from the previous studies (see above) and we obtained *An. gambiae* orthologs when needed using FlyBase (43), VectorBase (58) and OrthoMCL (77). We considered proximal enhancer-like regions those annotated by our approach to the same genes than the original targets in *Anopheles*, or any corresponding ortholog in *Drosophila*. To correct our annotation based on the nearest neighboring gene, we included the target genes identified by others when we could obtain an unambiguous single target gene from the published data sets. We considered a regulatory region to be potentially distal if located > 2 Kb away from the promoter of the target gene. To explore the relationship between the chromatin accessibility at these regions and expression of the annotated genes, we quantified the ATAC-seq signal at the THSs-overlapping enhancer-like regions as described above.

To discover novel TSS-like sites, *Drosophila* TSSs were downloaded from the Eukaryotic Promoter Database (Release 128-005, (78)), and as we did for enhancers, we used a homology-based approach to transfer coordinates to the *An. gambiae* genome. We then checked annotation to genomic features (HOMER), overlap with THSs (BEDTools *intersect*) and whether the annotated genes already have 5’ UTRs/TSS annotated, in order to classify these elements as novel.

To characterise CTCF binding sites, we downloaded binding sites for *Drosophila* CTCF from the ChIP-Atlas (79). As described above, we transferred the coordinates from *D. melanogaster* to the *An. gambiae* genome based in homology.

### Motif enrichment analysis

We performed d*e novo* motif analysis using HOMER. We applied this pipeline to different sets of THSs: activating or repressor, depending on the pattern of promoter accessibility and gene expression of the annotated gene. This analysis was conducted separately on the set of DiffBind regulatory regions at different locations and on the set of enhancer-like elements (see Results). We used the *findMotifsGenome*.*pl* module considering the THSs summit and searched for motifs in 100 bp in each direction (-size −100,100). See Supplementary Methods for additional details on this analysis.

## RESULTS

### Chromatin accessibility correlates with active transcription and epigenetic states

ATAC-seq libraries were produced from adult female *An. gambiae* s.s. mosquitoes infected with the *P. falciparum* parasite clone 3D7. Midguts (MGs) were dissected at 7 days post-infection and salivary glands (SGs) dissected at 14 days post-infection in two independent experimental infections (see Methods; Supplementary Table S1). After sequencing, the quality of the ATAC-seq data obtained was high and comparable between tissues and experimental infections (Figure 1A). Reproducibility analyses revealed a higher correlation of ATAC-seq between infections of the same tissue than between tissues, clustering by tissue in a Principal Components Analysis (PCA, Supplementary Figures S1A and B). Other quality control measurements, such as the fraction of reads mapping to the mitochondria, or the library complexity coefficients, also indicated the high quality of the ATAC-seq data according to ENCODE standards (Supplementary Table S2) (50). The fragment length distributions for both tissues conformed to previous observations (47,48); most insert sizes corresponded to nucleosome-free regions of less than 130 bp, and a second peak of fragment sizes represented mononucleosomes (Figure 1A). The distribution of the ATAC-seq nucleosome-free signal also matched the typical profiles of higher eukaryotes (53,80,81), with a higher density of insertions at the TSSs and two smaller peaks marking the spacing between adjacent nucleosomes (Figure 1B and Supplementary Figure S1C). By contrast, the highest density of ATAC-seq mononucleosome signal localized to positions flanking the TSSs at the +/-1 nucleosome positions (Figure 1B).

**FIGURE 1:**
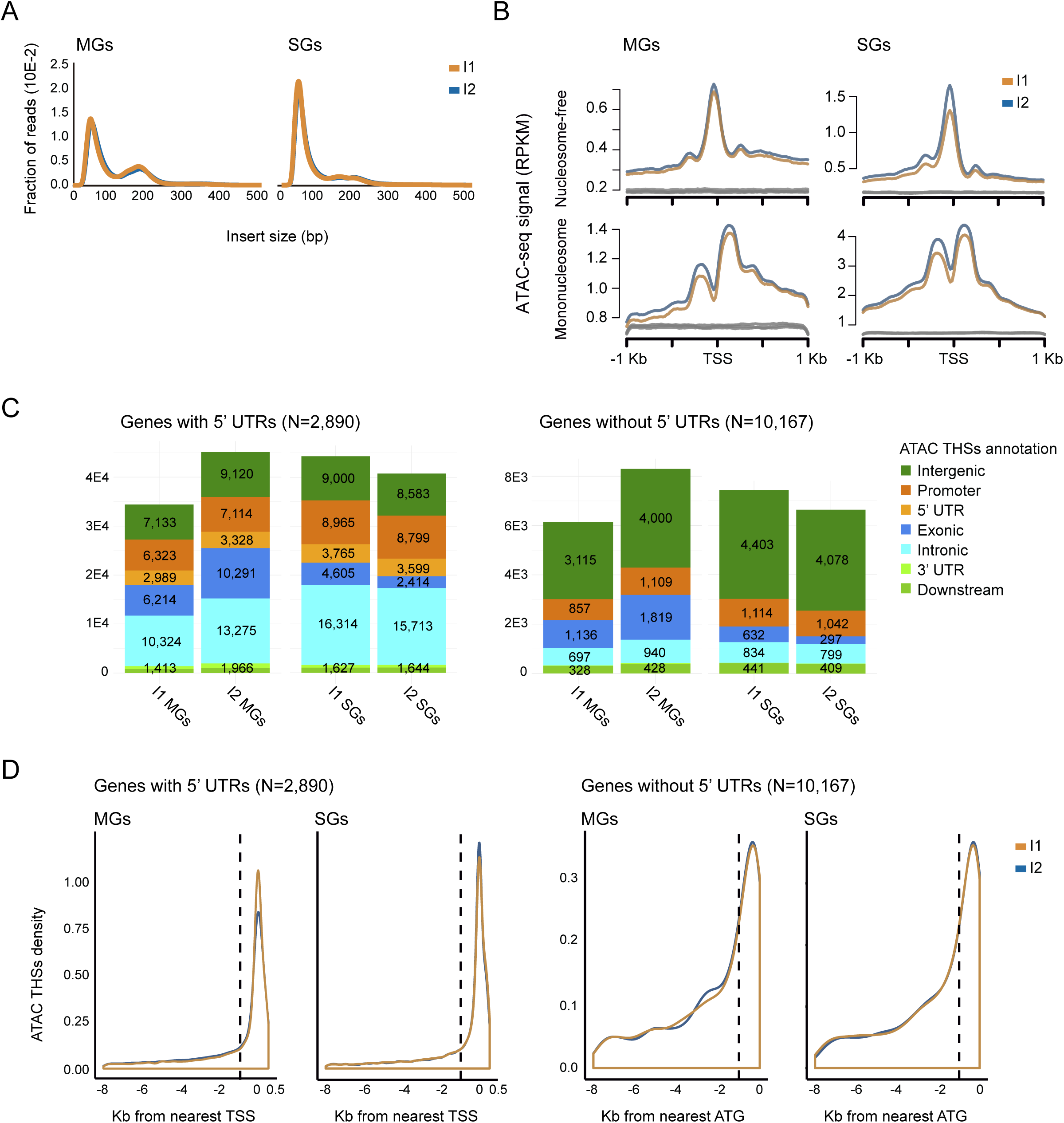
ATAC-seq allows for the genome-wide profiling of chromatin accessibility and nucleosome occupancy in *An. gambiae*. **(A)** ATAC-seq fragment size distribution corresponding to *P. falciparum*-infected *An. gambiae* midguts (MGs) and salivary glands (SGs). I1 and I2 are biological replicates (independent infections). A large proportion of reads are less than 100 bp, which represents the nucleosome-free region. The plot also shows a clear periodicity, which is indicative of nucleosome occupancy. To filter THSs (nucleosome-free regions) and mononucleosomes, we selected reads in the ranges of 50-130 bp and 171-256 bp, respectively. **(B)** Average profile plots of normalized (RPKM) ATAC-seq nucleosome-free and mononucleosomal reads surrounding *An. gambiae* annotated TSSs (±1 Kb). Higher mononucleosomal signals flank the nucleosome-free region at TSSs. Profiles in grey represent reads density at random genomic coordinates. **(C)** Annotation of THSs to features genome-wide: intergenic regions, promoters, 5’ UTRs, exons, introns, 3’ UTRs and downstream regions. Most THSs annotate to introns. **(D)** Density plot showing the position of THSs with respect to the TSSs (or ATG start codons for genes without annotated 5’ UTRs). Higher densities of THSs occur within 1 Kb upstream the TSSs or ATG sites. The dashed lines indicate the putative promoter region located 1 Kb upstream.

Once we had validated the ATAC-seq approach, we used the MACS2 peak-calling software (54) to identify regions significantly enriched in ATAC-seq nucleosome-free signal, which we denote Tn5 hypersensitive sites (THSs). The computed THSs for all samples are listed in Supplementary Table S4 and include a total of 111,586 unique accessible regulatory regions (43,010 in MGs and 68,576 in SGs) that were present in both experimental infections. The total number of THSs differed slightly between tissues (see Methods), but we observed that average ATAC-seq enrichment levels at THSs were similar across samples (Supplementary Figures S1D and E). These THSs annotated to > 10,000 genes (of a total of 13,057 genes in *An. gambiae*). Approximately 20% of the total *An. gambiae* genes do not have 5’ UTRs/TSS annotated, and the promoter for those genes is assumed to lie in the 1Kb region upstream of the ATG start codon (see Methods). The annotation of the THSs to genomic features showed a preferential location within introns (30.5%), followed by promoters (18.3%), exons (14.2%), and 5’ UTRs (7.1%) (Figure 1C and Supplementary Table S4). The concentration of accessible sites within introns has been previously reported, for example in *D. melanogaster* tissues, with around 50% of ATAC-seq THSs found within introns (82). The density of THSs at promoters diminishes with distance from the TSS. For genes without annotated 5’ UTRs, THSs density also decreases with distance from the ATG start codon of the gene (Figure 1D).

In addition to THSs, ATAC-seq data is also informative about patterns of nucleosome positioning. The pattern for metazoans is such that the 5’ promoter appears nucleosome-free and the transcribed regions are occupied by a more periodic array of positioned nucleosomes (83,84) Similarly, in our study we observed a larger proportion of mononucleosome signal at exons and introns compared to promoters and 5’ UTRs (Supplementary Figure S1F). To validate this more systematically, we used the NucleoATAC nucleosome calling software (63) to predict nucleosome dyads positions (symmetry axis of the nucleosomal DNA), and we found dyads within 100 bp windows of around 90% of the THSs (Supplementary Table S7).

Histone post-translational modifications (hPTMs) are known to play crucial roles in the remodeling of chromatin structure and transcriptional regulation, with *a priori* well-established activating (H3K9ac, H3K27ac, H3K4me3) or repressor (H3K9me3) functions (85,86). Activating hPTMs are expected to be enriched at mononucleosomes flanking active and accessible regulatory regions, like TSSs, whereas repressor histone marks are associated with non-accessible and nucleosome-occupied heterochromatin. To test such a relationship between chromatin accessibility and epigenetic states, we examined ATAC-seq signal relative to ChIP-seq peaks for various hPTMs that we obtained from our previous study on *P. falciparum*-infected and non-infected *An. gambiae* midguts (42). ATAC-seq nucleosome-free signals appeared more enriched (compared to a random distribution) at peaks of H3K9ac, H3K27ac, and to a lesser extent H3K4me3, and depleted at H3K9me3 peaks (Supplementary Figure S1G). Next, we applied the segmentation algorithm ChromHMM (71), that integrates the ATAC-seq and ChIP-seq data, to partition the genome. The purpose was to detect recurring epigenetic/accessibility patterns genome-wide and then assign a state to each region in the genome. This analysis resulted in eight chromatin states (Supplementary Figure S2A and Supplementary Table S8). Most of the genome (∼65%) appeared to be in a depleted state without ATAC-seq or ChIP-seq signal. Intergenic regions were largely depleted of ATAC-seq nucleosome-free signal and enriched in H3K4me3 and H3K9me3. By contrast, 5’ UTRs and promoters displayed states of open-chromatin, and activating hPTMs (H3K9ac, H3K27ac and/or H3K4me3). This was also the case for introns and exons, which also appeared in an open state (Figure 2A). Nearly half of the intronic THSs coincided with H3K27ac-enriched chromatin states (states 5/7; Supplementary Figure S2A) and only 7.5% with the chromatin state H3K9ac/H3K27ac/H3K4me3 (state 6; Supplementary Figure S2A and Supplementary Table S4). The observation of chromatin accessibility spanning into exons is also in agreement with the pattern previously reported in *Drosophila* (25,87).

**FIGURE 2:**
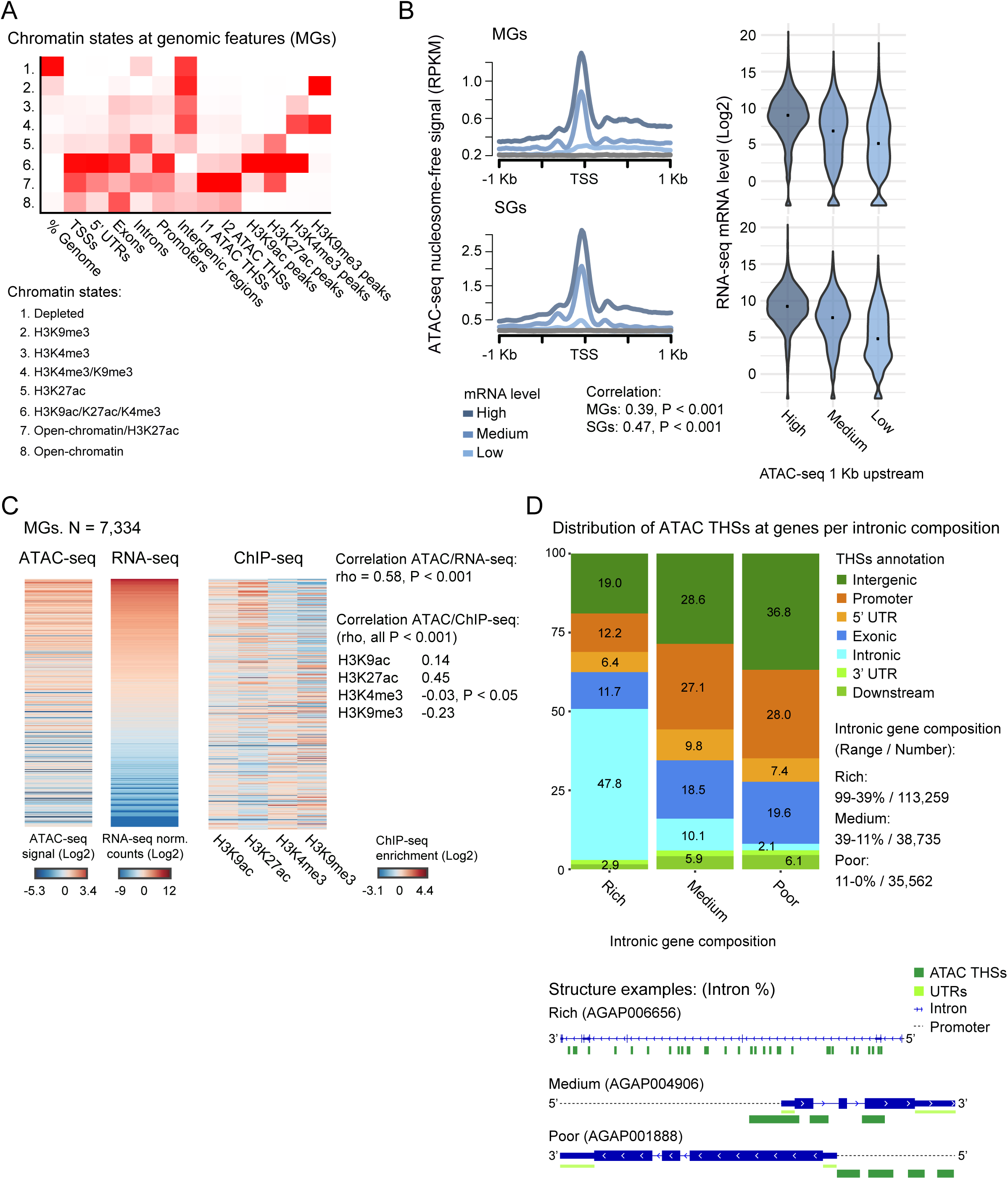
Chromatin accessibility by ATAC-seq is predictive of active hPTMs and tissue-specific gene expression. **(A)** Heatmap showing the overlap of various genomic features with chromatin states inferred using ChromHMM. Darker red in the first column indicates higher percentage of the genome overlapping with a particular state. In all other columns the red indicates the likelihood of finding a chromatin state compared to the random expectation. Most of the genome is in a depleted state. Introns and promoters display a typical state of open-chromatin and activating hPTMs. **(B)** Correlation between ATAC-seq nucleosome-free signal at TSSs and promoters and gene expression. Profile plots (left) show changes in ATAC-seq nucleosome-free signal enrichment at each tissue with respect to the TSSs (± 1 Kb). Genes are divided into groups and ranked by their mRNA levels (high, medium or low). Violin plots (right) show mRNA levels for genes grouped by their level of ATAC-seq nucleosome-free signal at promoters: high, medium and low chromatin accessibility. Plot width accounts for the density of repeated values in the range. Median values are marked with a black dot. **(C)** Heatmap showing ATAC-seq nucleosome-free enrichment, gene expression levels of the annotated gene, and hPTMs enrichment at promoters. Data correspond to a subset of non-overlapping genes with a THSs annotated in which there is a positive relationship between accessibility and gene expression, i.e. transcriptional activation (see Methods). The plot is for midguts. Genes are ordered by mRNA levels. ATAC-seq and ChIP-seq enrichments at promoters are normalized (RPKM) and the ChIP-seq input-corrected. Data are mean-centered. **(D)** Frequency of THSs annotated to various regions for genes grouped by their intronic composition: intron-rich, intron-medium and intron-poor. THSs tend to localise at introns, rather than at the promoter, in high intronic genes, and the opposite is true for low intronic genes. Diagrams show the archetypal structure for the three categories of genes based on their intronic composition.

The integration of ATAC-seq and RNA-seq data can be used to infer clusters of co-regulated genes and common regulatory mechanisms. Looking at the association between nucleosome occupancy and gene expression, mononucleosomes were positioned more frequently at the promoters of more highly-expressed genes (Supplementary Figure S2B and Supplementary Table S5). Our results also showed that nucleosome-free ATAC-seq enrichment at promoters was positively associated with gene expression levels of the corresponding genes (Spearman test: rho 0.39, P < 0.001 (MGs); rho 0.47, P < 0.001 (SGs); Figure 2B). To further investigate this pattern in a more quantitative manner, we filtered non-overlapping genes with THSs located at promoters or 5’ UTRs, and categorized them by their expression and chromatin accessibility levels at promoters (see Methods). In approximately 90% of cases, the high/medium expressed genes appeared in the high/medium promoter accessibility groups and conversely, medium/low expressed genes appeared in the medium/low promoter accessibility categories (Figure 2C; Supplementary Figure S2C and Supplementary Table S5). This suggests that the function of these regulatory regions is gene activation. Notably, around 10% of the THSs-annotated genes displayed opposite profiles for chromatin accessibility and gene expression levels (high promoter accessibility/low expression or low promoter accessibility/high expression; Spearman test: rho −0.71, P < 0.001 (MGs); rho −0.65, P < 0.001 (SGs); Supplementary Figure S2D and Supplementary Table S5), suggesting that these accessible THSs correspond to binding sites for repressor TFs. Finally, by integrating ATAC-seq and RNA-seq data with ChIP-seq data, we observed that higher accessibility and expression correlated positively with H3K9ac/H3K27ac enrichment in the promoter, and negatively with H3K9me3 in MGs (Spearman test: rho 0.14, P < 0.001 (H3K9ac); rho 0.45, P < 0.001 (H3K27ac); rho −0.23, P < 0.001 (H3K9me3); Figure 2C).

A large proportion of THSs (30.5%) were located within introns, suggesting that accessibility in these regions may have a role in the regulation of gene expression. Consistent with this hypothesis, genes that are more intronic relative to the gene length (intron-rich genes, see Methods) were found to have a higher number of THSs that accumulate within introns. Conversely, for intron-poor genes, the THSs tended to be located at promoters (Figure 2D). This pattern is independent on whether the genes have 5’ UTRs annotated or not (genes with annotated 5’ UTRs: intron-rich 49.3% intronic/12.5% promoter THSs, intron-poor 2.3% intronic/31.8% promoter THSs; genes without annotated 5’ UTRs: intron-rich 31.2% intronic/8.9% promoter THSs, intron-poor 1.3% intronic/16.3% promoter THSs). Indeed, genes without annotated 5’ UTRs did not seem to accumulate more THSs within introns when compared to genes with annotated 5’ UTRs (Mann-Whitney U test: P > 0.05; Figure 1C). The mean intronic accessibility was positively correlated with chromatin accessibility at promoters (Spearman test: rho 0.33, P < 0.001 (MGs); rho 0.41, P < 0.001 (SGs)). There was also a positive and significant correlation between mean intronic accessibility and gene expression (Spearman test: rho 0.20, P < 0.001 (MGs); rho 0.27, P < 0.001 (SGs)), which was higher for intron-rich genes (rho 0.25, P < 0.001 (MGs); rho 0.33, P < 0.001 (SGs)), and when considering the first intron only (rho 0.32, P < 0.001 (MGs); rho 0.45, P < 0.001 (SGs)). Quantitatively, we also observed that intron-rich genes tend to be more expressed (Spearman test: rho 0.16, P < 0.001 (MGs); rho 0.30, P < 0.001 (SGs); Supplementary Figure S2E)). Overall, these observations suggest that chromatin accessibility within introns could be functionally involved in gene expression regulation and is dependent on the gene architecture.

Among the target genes with a THS annotated, we found 241 important immune-related genes such as *stat2, rel2* and members of the *TEP* family (62) (Supplementary Table S4), as well as 671 genes that we reported in an earlier study as *Plasmodium*-responsive by comparing *P. falciparum*-infected and non-infected *An. gambiae* midguts (42). These include, for example, genes encoding CLIP serine proteases, argonaute 1 and the defensin DEF1 (Supplementary Table S4). Regulatory sites at these genes are located predominantly within introns (39.4%), but also at 5’ UTRs (7.2%) and promoters (11.5%). Lastly, for this set of genes we also report here a significant and positive correlation between chromatin accessibility and gene expression levels (MG, Spearman test: rho 0.31 P < 0.001 (promoters); rho 0.10, P < 0.05 (introns); Supplementary Figure S2F).

Overall, our results show a genome-wide correlation between chromatin accessibility at regulatory regions (i.e. promoters and introns) and the gene expression levels for each tissue assayed. We also confirm that these regulatory sites display the typical pattern of histone post-translational modifications (hPTMs) characteristic of active chromatin.

### Tissue-specific chromatin accessibility correlates with differential gene expression

We found evidence for tissue-specific chromatin accessibility profiles through differential chromatin accessibility analyses at THSs between *An. gambiae* MGs and SGs. A higher proportion of differentially accessible regions (DiffBind regions) were more accessible in SGs (85.1%, n=21,243) compared to MGs (Supplementary Figure S3A and Supplementary Table S6), and the majority were located at promoters (26.9%) or within introns (30.2%) (Supplementary Figure S3B and Supplementary Table S6). The majority of the DiffBind regions corresponded to changes in the level of accessibility between tissues, rather than a presence/absence of a THS in one of the tissues (Supplementary Figure S3C). In total, 21,400 DiffBind regions coincided with a THS present in both biological replicates, and we used this high-confidence set for downstream analysis (Supplementary Table S6).

The integration of the ATAC-seq and RNA-seq data for genes that appeared differentially expressed and accessible between the two tissues (1,920 out of a total of 3,584 DEGs; Supplementary Figure S3D and Supplementary Table S6), revealed a significant correlation between the level of accessibility at the regulatory sites and the levels of gene expression in the same tissue (5,382 Diffbind regions at 5’ UTRs, promoters or introns of 1,920 DEGs; Spearman test, rho 0.25, P < 0.001 (MGs); rho 0.12, P < 0.001 (SGs); Figure 3A). Additionally, there was also a positive correlation between the magnitude of the change in accessibility at the DiffBind regions within regulatory sites of DEGs and the change in gene expression between tissues (Spearman test, promoters: rho 0.37, P < 0.001; introns: rho 0.19, P < 0.001; Figure 3B). Full details of these analyses are given in Supplementary Info.

**FIGURE 3:**
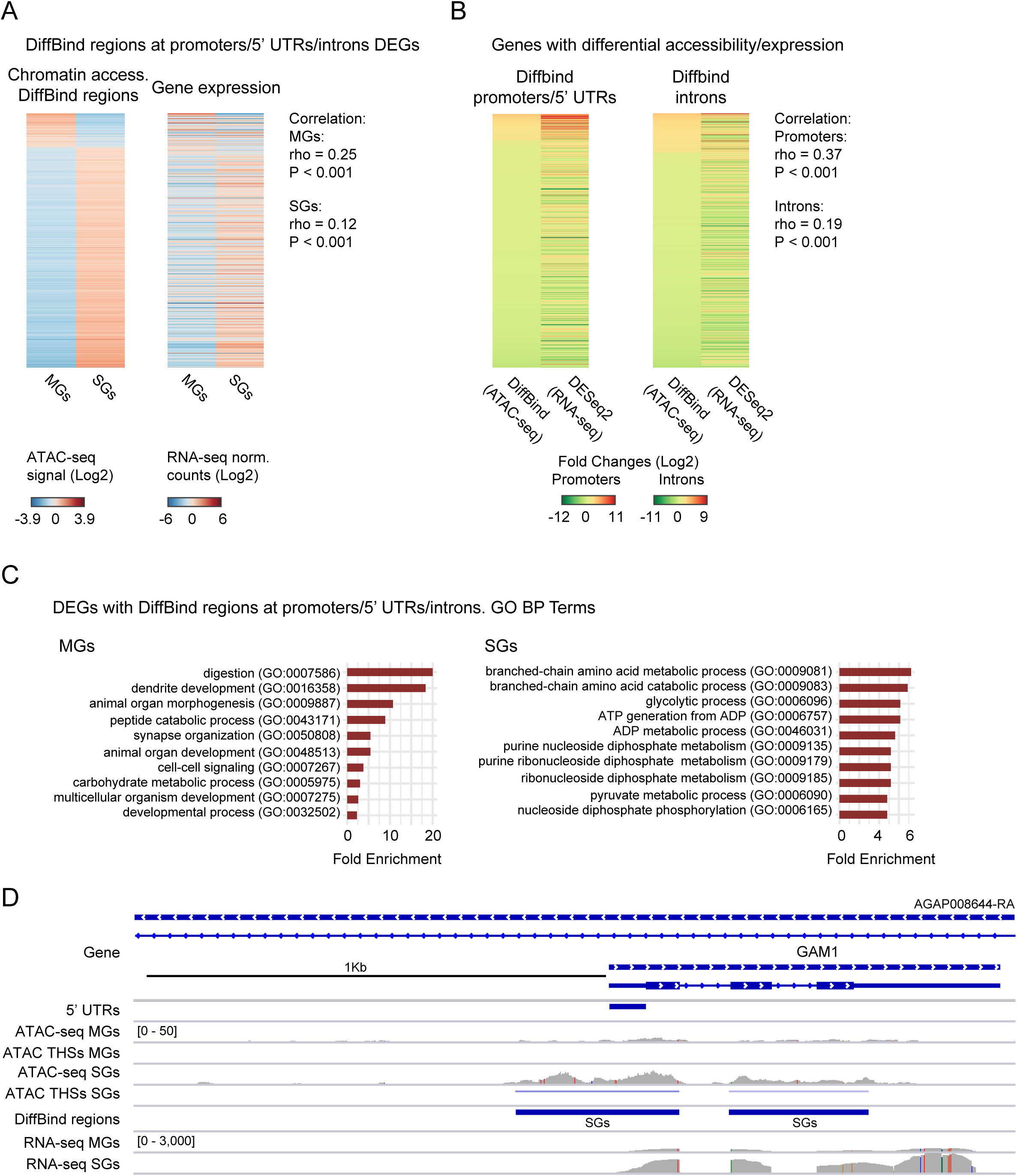
Differential chromatin accessibility between tissues correlates with changes in gene expression. **(A)** Heatmap showing ATAC-seq nucleosome-free enrichment at DiffBind regions located at 5’ UTRs, promoters or introns of differentially expressed genes and their expression levels. There is a positive and significant correlation between chromatin accessibility at these regions and gene expression. Genes are ordered by normalized ATAC-seq enrichment (RPKM). Data are mean-centered and for infection 2. **(B)** Heatmap showing chromatin accessibility and gene expression fold changes between tissues for differentially expressed genes that display a DiffBind region at the promoters and/or 5’ UTRs and at the introns. Changes occur in most cases in the same direction and there is a positive and significant correlation between the magnitude of changes in accessibility and expression. **(C)** Top GO Biological Processes terms overrepresented in the set of DEGs with DiffBind regions located at promoters, 5’ UTRs and/or introns for each tissue. **(D)** Chromatin accessibility and gene expression profiles in the region containing the antimicrobial peptide GAM1-encoding gene (AGAP008645) which is differentially expressed and differentially accessible between tissues. The tracks displayed are for midguts and salivary glands from infection 2. The location of 5’ UTRs, THSs, and the regions of differential accessibility (DiffBind) are indicated by colored bars. All tracks are shown at equal scale.

The GO overrepresentation functional analyses, performed on the DiffBind regions at 5’ UTRs, promoters or introns of DEGs, showed different processes to be enriched in each tissue, correlating with different functional activities such as digestion and peptide catabolism in the MGs, or amino acid metabolism and glycolysis in the SGs (Figure 3C; Supplementary Figure S3E and Supplementary Table S6). These results conform with the processes expected for each tissue. For example, genes related to digestion, such as trypsins, are known to be specifically upregulated in a tissue-specific manner in MGs after a blood meal (88). Previous studies have also shown the upregulation of glycolytic processes in mosquito SGs (89,90) and of amino acid metabolism in *Anopheles* SGs in response to *Plasmodium* infection (91). In addition, we found more expression linked with more accessible promoters in the same tissue for immune-related genes, such as proteases in MGs and serpins in SGs (Supplementary Figure S3E). These included, for example, serpins involved in the Toll pathway (SRPN2 and SRPN6), and which have been linked to the *An. gambiae* immune response to *P. berghei* sporozoites in the SGs (92). Other examples of mosquito genes in which a functional link was established are TF-encoding genes, such as *rel2*, which modulates anti-*Plasmodium* factors, and other immune-related genes including the C-type lectin *ctl4* and the defensin *def1* (Supplementary Table S6). The gene encoding the gambicin antimicrobial peptide *gam1* (AGAP008645) is a good case example. This gene was more expressed in SGs, and this upregulation coincides with several regions of higher chromatin accessibility located at the TSS/promoter and spanning into exons and introns (Figure 3D).

A high proportion of the THSs and DiffBind regions were found within introns (44.7%) or exons (36.6%), so they could be also implicated in regulation of expression at the isoform rather than the gene level. Our analysis revealed evidence of isoform switching at 176 *An. gambiae* genes, with 346 isoforms changing between the two infected tissues (Supplementary Figure S3F and Supplementary Table S9). The majority (90%) of these genes contained DiffBind regions, mostly within introns (61.7%) or at promoters (24.6%) (Supplementary Figure S3G). These results suggest a functional link between chromatin accessibility dynamics at regulatory regions (mainly introns) and gene isoform switching.

We performed motif enrichment analysis on the regions with differential chromatin accessibility to predict the sets of TFs that may be involved in tissue-specific functional gene expression. By comparing the tissue in which the regulatory sites were more accessible and the annotated genes more expressed, we could infer the functions of the TFs in transcriptional activation or repression (i.e. higher accessibility in a tissue corresponded to higher or lower expression in the same tissue; Supplementary Table S6). We first focused on the set of DiffBind regions annotated to 5’ UTRs or promoters for DEGs specific to midguts (MGs) and salivary glands (SGs) (Supplementary Table S10). For the subset of accessible regions that annotate to active genes, we found *de novo* motifs that match consensus sequences (binding sites) for TFs that are known *Drosophila* activators with midgut and salivary gland-specific functions such as serpent or odd paired (MGs) and homothorax or broad (SGs). The subset of regions in which the accessibility change was linked to silencing were enriched in motifs similar to binding sites of known *Drosophila* repressors, such as even, skipped or pleiohomeotic for midguts, and forkhead or hunchback specific to salivary glands. Next, provided that the majority of DiffBind regions were located within introns (see above) we performed analogous analyses to predict TFs at these regions (Supplementary Table S10). In the majority of cases, we found enriched motifs similar to the ones in promoters and 5’ UTRs (see above). We also predicted TFs that appeared to be particular to introns, such as bric a brac 1 or schnurri.

Altogether, chromatin accessibility and gene expression differential analyses between tissues allowed us to identify mosquito genes in which the switch between the open/closed chromatin is associated with transcriptional activation in a tissue-specific manner. We also provide some evidence that differential accessibility at intronic regulatory regions could be related with changes in isoform rather than gene expression. Lastly, the motif enrichment analyses allowed us to predict binding sites for potential activator or repressor TFs with tissue-specific functions.

### Chromatin accessibility allows for the identification of novel *An. gambiae* cis*-*regulatory elements

ATAC-seq has been shown to capture regulatory sequences with high precision, and therefore is an ideal assay to characterize novel cis-regulatory elements such as enhancers and TSSs. Of the 1,685 *An. gambiae* enhancers suggested from previous studies (37,38,44), mostly based on computational predictions, 42% (708) were identified as THSs (Supplementary Table S11), and therefore may correspond to active enhancers in the tissues assayed here. Furthermore, using a homology approach and *D. melanogaster* enhancer maps such as the EnhancerAtlas (27) (see Methods and Supplementary Info), we predicted 1,122 novel enhancer-like regions. Of these regions, around 10% (103) were found to overlap with THSs by ATAC-seq (Supplementary Table S11). As a consequence, the final database comprises a total of 811 accessible enhancer-like regions in *An. gambiae*, overlapping with 2,272 THSs annotated to 563 genes. The majority located within introns (43.3%) or promoters (21%) (Figure 4A and Supplementary Table S11). Of these 811 accessible enhancer-like regions, 293 displayed signatures typical of active enhancers: chromatin accessibility and H3K27ac enrichment (Figure 4B and Supplementary Table S11). For around 80% (633) of the accessible enhancer-like regions defined by ATAC-seq that we annotated here to the closest gene, that gene coincides with the target of the enhancer reported previously by others (26,38,44) and therefore these are most likely to be proximal enhancer elements. The remainder (178) may be distal enhancers (see Methods and Supplementary Info; Supplementary Table S11). There was a significant association between chromatin accessibility at the regulatory sites and the expression of the annotated target genes for proximal enhancers (Spearman test: rho 0.35, P < (MGs); rho 0.15, P < 0.05 (SGs); Figure 4C), but not for distal (P=0.66 for MGs and P=0.32 for SGs).

**FIGURE 4:**
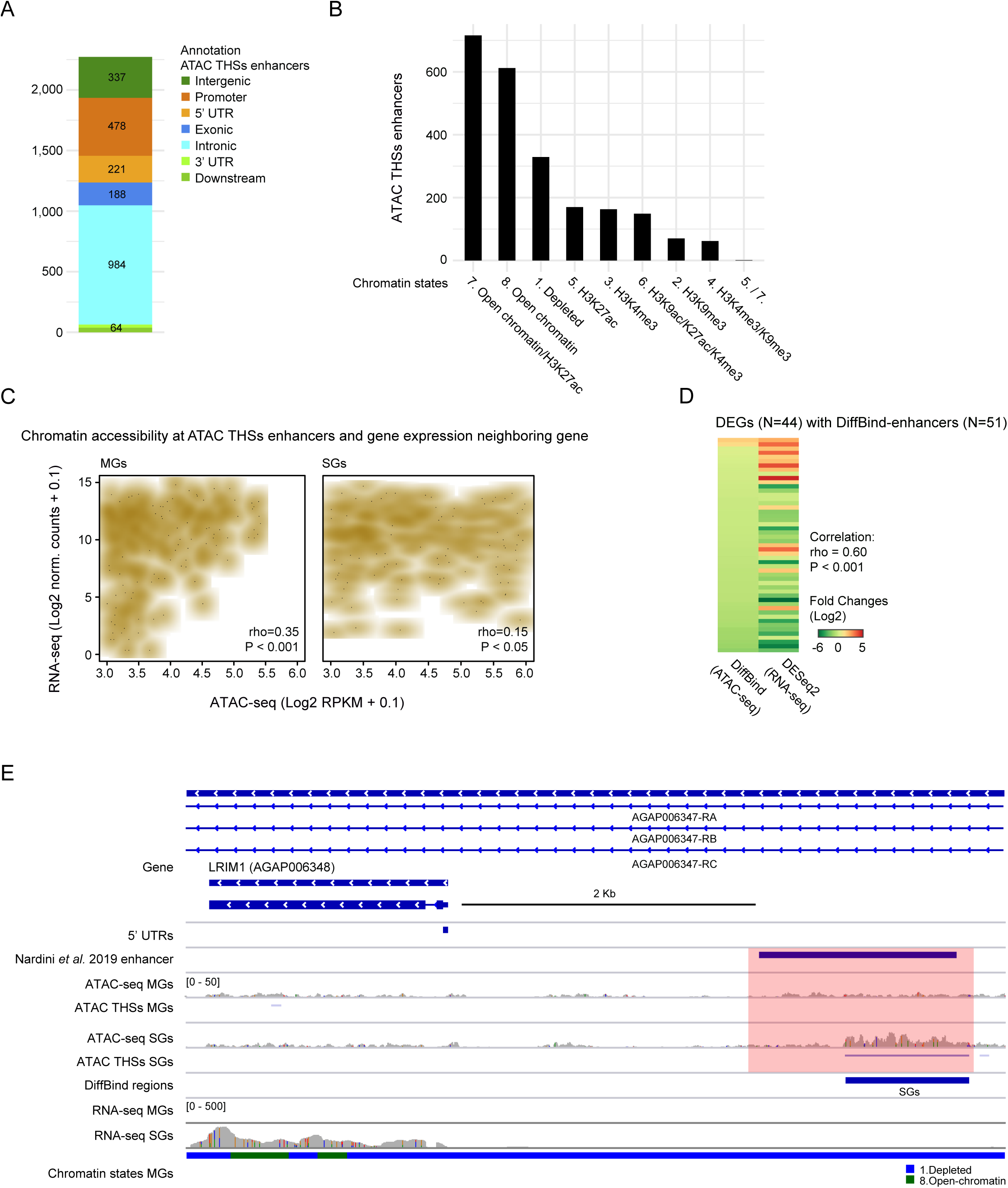
Genome-wide *in vivo* mapping and functional characterisation of *An. Gambiae* enhancers. **(A)** Annotation of accessible enhancers to various genomic features: intergenic regions, promoters, 5’ UTRs, exons, introns, 3’ UTRs and downstream regions. The majority of enhancers locate at introns or promoters. **(B)** Chromatin states at the accessible enhancers. As expected, these regulatory regions appear to be H3K27ac-enriched. **(C)** Scatter plots showing a positive correlation between chromatin accessibility for a subset of proximal enhancers (see Results), and gene expression of the nearest target gene. Data is for midguts (left) and salivary glands (right). **(D)** Heatmap displaying chromatin accessibility and gene expression fold changes between tissues for differentially expressed genes that show a DiffBind region coinciding with a proximal enhancer element. Changes occur in most cases in the same direction and there is a positive correlation in the magnitude of the change. **(E)** Chromatin accessibility and gene expression profiles in the region containing the LRIM1-encoding gene (AGAP006348), a *Plasmodium*-responsive gene based on our previous study (42). Here, this gene is differentially expressed and displays a differentially accessible enhancer between tissues. The tracks shown are for midguts and salivary glands from infection 2. All tracks are shown at equal scale. The location of various genomic features: 5’ UTRs, THSs, DiffBind regions and chromatin states, are indicated by colored bars. The enhancer element as predicted by others (44) that is coinciding with the differential accessibility region is highlighted as the pink box.

The integration of the differential gene expression data between the tissues and differential chromatin accessibility at enhancer-like regions was used as a proxy to validate functions. 51 regions were identified both as proximal enhancer-like regions and DiffBind regions, and annotated to differentially expressed genes between the two tissues. Of these, the most frequent role was activating: in 37 cases the enhancer region was more accessible in the tissue where the gene was more highly expressed. However, in 14 cases the relationship was the opposite, pointing to a repressor function. There was also a significant positive correlation between the changes in accessibility and the differential expression of the annotated genes (Spearman test: rho 0.60, P < 0.001) (Figure 4D). The *lrim1* gene (AGAP006348), that encodes for a leucine-rich immune protein, is a good example of a *Plasmodium*-responsive gene (42), that has an experimentally validated enhancer and that was differentially accessible between tissues. Here, we observed the gene to be more highly expressed in SGs, and this was associated with a DiffBind region upstream of the promoter, which was more accessible in SGs, and that coincided with the enhancer experimentally validated by others (44) (Figure 4E). Apart from differences in gene expression, these regulatory elements may be also involved in differences in gene isoform expression. Indeed, there are 38 enhancer-like regions annotated to 19 genes with isoforms that switch expression. In addition, 20 out of these 38 enhancer-like regions contained instances of motifs that have been shown to be characteristic of *Drosophila* intronic-splicing enhancers, such as CTCTCT and TTATAA (93).

Finally, on the set of 2,272 THSs that overlap *An. gambiae* enhancer-like regions, we performed motif enrichment analyses to predict enriched *de novo* motifs similar to known binding sites of *Drosophila* TFs that could be acting through binding of enhancers. Among the top hits, we predicted TFs involved in processes such as nucleosome organization, reproduction and regulation of immune response (Supplementary Table S12). These included regulators such as trithorax-like, pleiohomeotic, zeste and the deformed epidermal autoregulatory factor-1, which had motifs that annotate to 306 genes.

ATAC-seq peaks can also be used to support TSS prediction and discovery. The majority of *An. gambiae* genes displayed THSs located at the promoter or the 5’ UTRs. In 46.5% of those, the THSs coincided with the annotated TSSs (i.e. the 5’ coordinate of the annotated UTR). Around 20% of genes in the current *An. gambiae* genome annotation (2,890) do not have annotated 5’ UTRs. We observed that 35% of these (1,009) displayed THSs located at promoters, which could be novel TSSs (Supplementary Table S4), and in agreement, 51% of them displayed chromatin states characteristic of TSSs (i.e. open-chromatin, H3K9ac, H3K27ac, and/or H3K4me3 enrichment) (Supplementary Table S11). To validate novel TSSs, and based on the assumption that TSSs tend to be conserved (94,95), we used a *Drosophila* dataset (78) and identified 917 homolog TSS-like sites in *An. gambiae* annotated to 819 genes (Supplementary Table S11). Integrating this set of homolog TSS sites with the mosquito ATAC-seq data, we found that the 28.5% of the transferred TSSs (217 out of 917) overlapped with our THSs at promoters or 5’ UTRs (Supplementary Figure S4A), and displayed enrichment in active hPTMs characteristic of promoter regions (Supplementary Figure S4B). As a proof of principle, 79.7% (173) of homolog TSSs validated mosquito annotated TSSs, and for the rest, the THSs did not coincide with the annotated TSSs and thus could be considered as alternative TSSs. Finally, we report 14 potentially novel TSSs that annotated to genes without 5’ UTRs (Supplementary Table S11).

A third type of cis-regulatory sequences are insulators, specific DNA sequences that are bound by insulator binding proteins (IBPs) and play an important role in regulating gene expression (96,97). The CCCTC binding factor (CTCF) is a transcription factor known to bind insulators and domain boundaries in vertebrates and *Drosophila*. It contributes to long-range chromatin interactions, including enhancer-promoters, and organization of chromatin architecture. In *An. gambiae* the CTCF-like gene (AGAP005555) is the ortholog to CTCF in *Drosophila* and appears to be expressed in the tissues assayed here. Based on the assumption that the binding sites for CTCF as determined by ChIP-seq in *Drosophila* (79) should be highly conserved in *Anopheles* (98), we could identify 2,516 homolog *An. gambiae* CTCF sites, which we propose could function as mosquito insulator elements. 30.4% of the transferred CTCF sites (764 out of 2,516) overlapped with our ATAC-seq THSs (Supplementary Table S11). For 51.3% of these (392 out of 764), the annotated gene was ortholog to the nearest neighboring gene to the CTCF binding sites in *Drosophila*. We also report a fraction of potential CTCF binding sites that coincide with differentially accessible regions between midguts and salivary glands (177 out of 2,519). Of these, 29 were annotated to differentially expressed genes such as the ortholog gene in *An. gambiae* of the abdominal A (abd-A) gene in *Drosophila* (Supplementary Figure S4C), which is part of the Bithorax Complex (97).

## DISCUSSION

The aims of this study were to investigate mechanisms underlying tissue-specific regulation of gene expression, and to map genome-wide enhancer- and TSS-like activity in *An. gambiae in vivo*. In a previous study, we characterized different post-translational modifications of histone tails (hPTMs), and the transcriptional profiles of *P. falciparum*-infected and non-infected *An. gambiae* midguts, which allowed us to identify changes in the epigenomic landscape of the mosquito linked to malaria infection (42). However, this information was insufficient to capture with high precision the location and function of tissue-specific mosquito cis-regulatory elements (CREs). To this end, in this study we performed the first genome-wide chromatin accessibility profiling by ATAC-seq, together with gene expression analysis by RNA-seq, in *An. gambiae* tissues infected with *P. falciparum*.

We report thousands of accessible regulatory sequences (THSs) involved in tissue-specific transcriptional regulation, which were distributed genome-wide, particularly at introns and promoters (regions 1 Kb upstream of genes). This pattern is in agreement with the distribution of accessible regulatory sites reported in *Drosophila* by ATAC-seq (25,82), and in *Ae. aegypti* and *An. gambiae* mosquitoes by FAIRE-seq (39-41). Chromatin accessibility at regulatory sites is generally considered a good predictor of gene activity (50,99), which our results support, showing a positive correlation between open-chromatin at regulatory regions and gene expression. By integrating our ATAC-seq and ChIP-seq data for various hPTMs (42), we also describe a relationship between accessibility and epigenetic states at these regulatory sites: the enrichment of hPTMs with a priori activating (H3K9ac, H3K27ac) or repressor (H3K9me3) roles that relate to gene function.

In *D. melanogaster*, introns are known to harbour regulatory sequences and to have an important role in the spatio-temporal control of gene expression (100,101). In *An. gambiae*, we demonstrate a relationship between accessibility at introns, gene expression, and H3K27ac enrichment at the active site, suggesting that intronic THSs are involved in functional regulation of gene expression. Another important observation is that gene architecture influences the proportion of open CREs at introns respective to promoters, and vice versa: genes with higher intronic content contain more intronic THSs, and moreover, chromatin accessibility at these regions also correlates with higher accessibility of the cognate promoters and higher gene expression. Our results agree with previous hypotheses that longer introns would be more efficient in transcriptional enhancement (102), and is also in agreement with observations that introns can influence transcription by looping or coupling with promoters, transcribing sRNAs, or accumulating regulatory sequences (8,101). This is a poorly understood phenomenon that has been termed intron-mediated enhancement. It seems to be conserved among eukaryotes, including *Drosophila* (103,104).

Our comparative analyses of mosquito midguts and salivary glands allowed us to unveil tissue-specific regulatory elements that may underlie functional differences between tissues. We identified thousands of THSs displaying differential chromatin accessibility between tissues, annotated to differentially expressed genes. A major proportion of differentially-accessible regions were more accessible in salivary glands, and were located at promoters or introns. A higher fraction of ATAC-seq nucleosome-free reads were seen at salivary glands when compared to midguts, which could reflect both that the ATAC-seq assay worked better in this tissue, or that salivary glands display a more accessible regulatory landscape. Notably, the integration of our differential datasets revealed a correlation between accessibility and the transcriptional state, i.e. genes tended to be more expressed in the tissue where the regulatory sites were more accessible. Moreover, these genes appeared to have tissue-specific functions, such as digestion in midguts and amino acid metabolism in salivary glands, which has also been shown to be a pathway affected by *Plasmodium* infection (91). The next step was to predict the regulatory proteins involved in these functional responses. Here we report that the CREs with tissue-specific accessibility appear enriched in motifs resembling the consensus binding sites of *Drosophila* TFs involved in tissue-specific processes, including TFs with immune functions in midguts and salivary glands such as srp and rel. The functions of most of these predicted *Drosophila* TFs are likely conserved in mosquitoes (105,106). Indeed, many of the TFs predicted in this study, such as srp, Deaf1 or Trl, agree with those predicted by previous studies that employed FAIRE-seq in different *Ae. aegypti* and *An. gambiae* tissues (39,40). Among the differentially-expressed genes that seem to be regulated by differentially accessible CREs and that contain the above motifs, we found examples of immune-related genes (62), such as *srpn6* or *gam1*, and genes that we identified in our previous study as *Plasmodium*-responsive (42), such as the defensin *def1*. The regulation of these genes is likely to be crucial in determining mosquito infection phenotypes that impact traits such as vector competence, immune response, longevity or reproduction. Until now, the regulatory elements of most immune-related and *Plasmodium*-responsive *An. gambiae* genes have been uncharacterized.

Another motivation of this study is the genome-wide mapping, discovery and validation of functional enhancers and TSSs in *An. gambiae;* several enhancer elements have previously been predicted bioinformatically, but their *in vivo* characterization and functions remain poorly explored. Relatively few mosquito candidate enhancer sequences have been defined, compared to the four thousand enhancers described in *Drosophila* (EnhancerAtlas database, (27)), and only a very small number have been experimentally validated in *Anopheles* (38,44). Using our ATAC-seq data, we map *in vivo* 42% (708) of those 1,685 *An. gambiae* enhancers predicted bioinformatically (37,38,44). In addition, we found 1,122 potentially novel *An. gambiae* regulatory regions that are homolog to known *D. melanogaster* enhancers (26), and which do not coincide with previously predicted mosquito regulatory sequences. Of these 1,122 enhancer-like regions, around 10% (103) were found to be accessible, by ATAC-seq analysis. The remaining 1,019 regions homolog to *Drosophila* enhancers were not accessible according to ATAC-seq analysis. This could be due to the fact that these enhancers were originally identified in *D. melanogaster* under different experimental conditions and tissues. In total we report 811 enhancer-like regions accessible according to ATAC-seq analysis, that are located throughout the genome, mainly in introns and exons. Our results also show that the majority of enhancer sites (around 80%) would regulate the neighboring gene. This distribution and pattern is in agreement with previous observations in other model organisms, which suggest enhancers are mainly proximal to or intragenic of target genes (26,107).

In support of the functional role of active enhancer sites in *An. gambiae*, a positive correlation was seen for the majority of enhancer sites between chromatin accessibility at proximal regulatory sites, and the gene expression of the target genes. A small number of potentially distal regulatory networks, which are known to play important roles during development and differentiation processes in *Drosophila*, were also identified in *Anopheles*, based on the location of ortholog genes in *Drosophila*.

Finally, a powerful tool to validate the enhancer function is the combined analysis of differential chromatin accessibility at enhancers and gene expression changes between tissues. We report 51 enhancer-like regions with differential chromatin accessibility in midguts and salivary glands, that annotated to differentially expressed genes. For 37 out of the 51, the enhancer was more accessible in the same tissue in which the gene was more expressed, and this association was also quantitative. Importantly, we observed that 141 of the accessible enhancers were annotated to genes that we previously reported to be *Plasmodium*-responsive (42). One example is the leucine-rich repeat protein 1 (LRIM1) encoding gene, that harbours an enhancer that switches chromatin states between tissues, and thus is implicated in tissue-specific gene expression regulation. Future studies in *An. gambiae*, applying approaches such as STARR-seq, transgenesis or Hi-C are now needed to validate the enhancers function.

In summary, we applied for the first time high-throughput genome-wide chromatin accessibility profiling by ATAC-seq in *Plasmodium*-infected *An. gambiae*, providing new evidence on the mechanisms of transcriptional regulation in mosquitoes. The integrative analyses of ATAC-seq, RNA-seq and ChIP-seq allowed us to link chromatin accessibility and structure with function, as well as to characterize tissue-specific CREs potentially involved in mosquito immune responses to *Plasmodium*. We also show ATAC-seq is a powerful tool for the *in vivo* discovery and characterisation of functionally active enhancers as well as insulator sequences, which are still poorly understood in mosquitoes. Such a detailed map of the regulatory genome of the main human malaria vector *An. gambiae* fills an important gap in the field, and it is essential for designing new strategies of disease control based on the genetic manipulation of mosquitoes.

## DATA AVAILABILITY

The ATAC-seq and RNA-seq data generated and analysed during the current study are available in the GEO repository under accession number GSE152924, https://www.ncbi.nlm.nih.gov/geo/query/acc.cgi?acc=GSE152924. Reviewers can access this information using the token slwpoccmdhibbmr. The datasets supporting the conclusions of this article are included within the article and its additional files.

## SUPPLEMENTARY DATA

Supplementary Data are available at NAR online.

## ACKNOWLEDGMENT

We thank Rodrigo G. Arzate-Mejía for edits on the manuscript and useful discussion. We are grateful to A Hidalgo-Galiana for her contribution in the technical ATAC-seq work, and to Dorothy Armstrong and Elizabeth Peat for their maintenance of the mosquito colonies and support for parasite cultures at the University of Glasgow.

