## Supplementary Material for "The regulatory genome of the malaria vector *Anopheles gambiae*: integrating chromatin accessibility and gene expression"

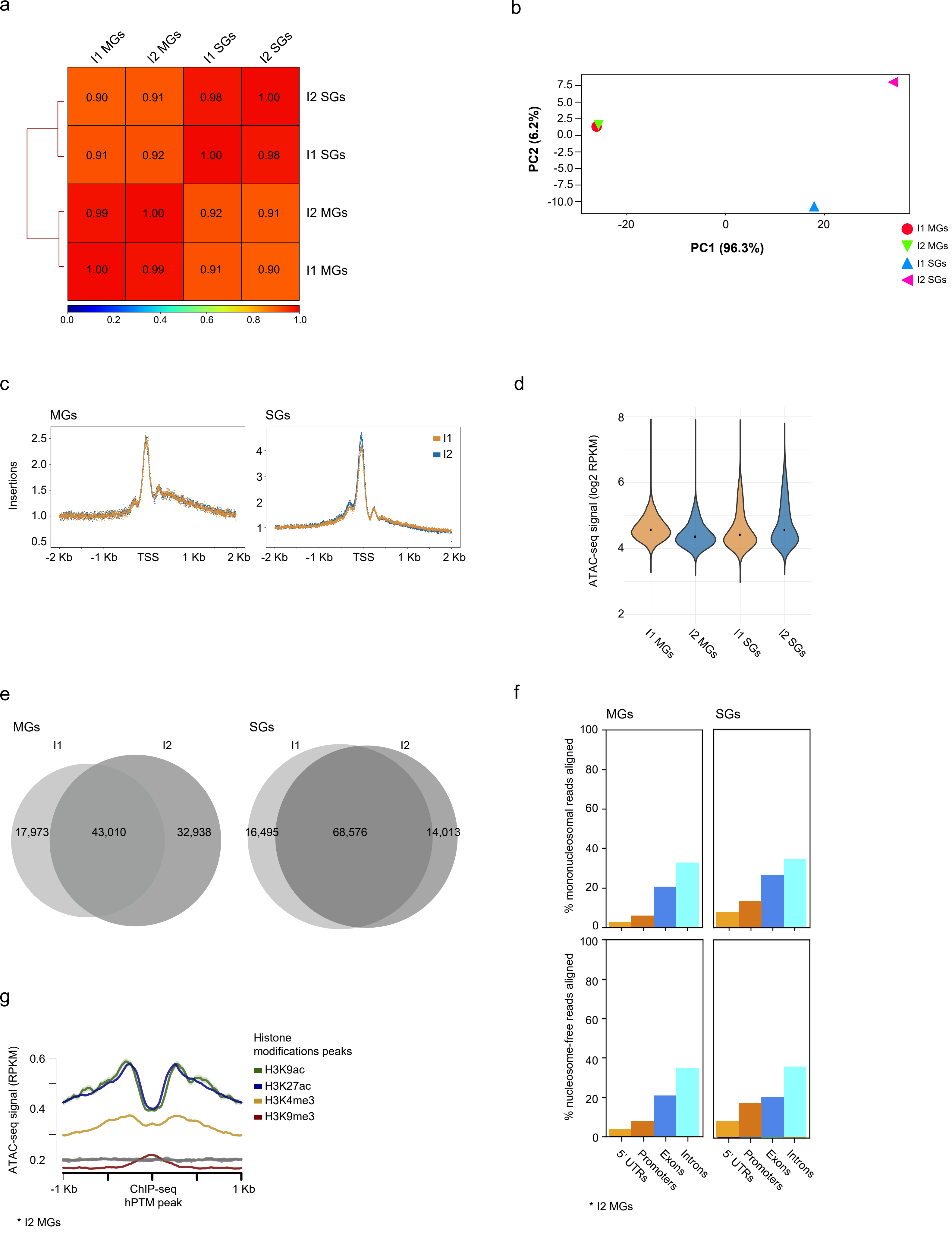


Supplementary Figure 1

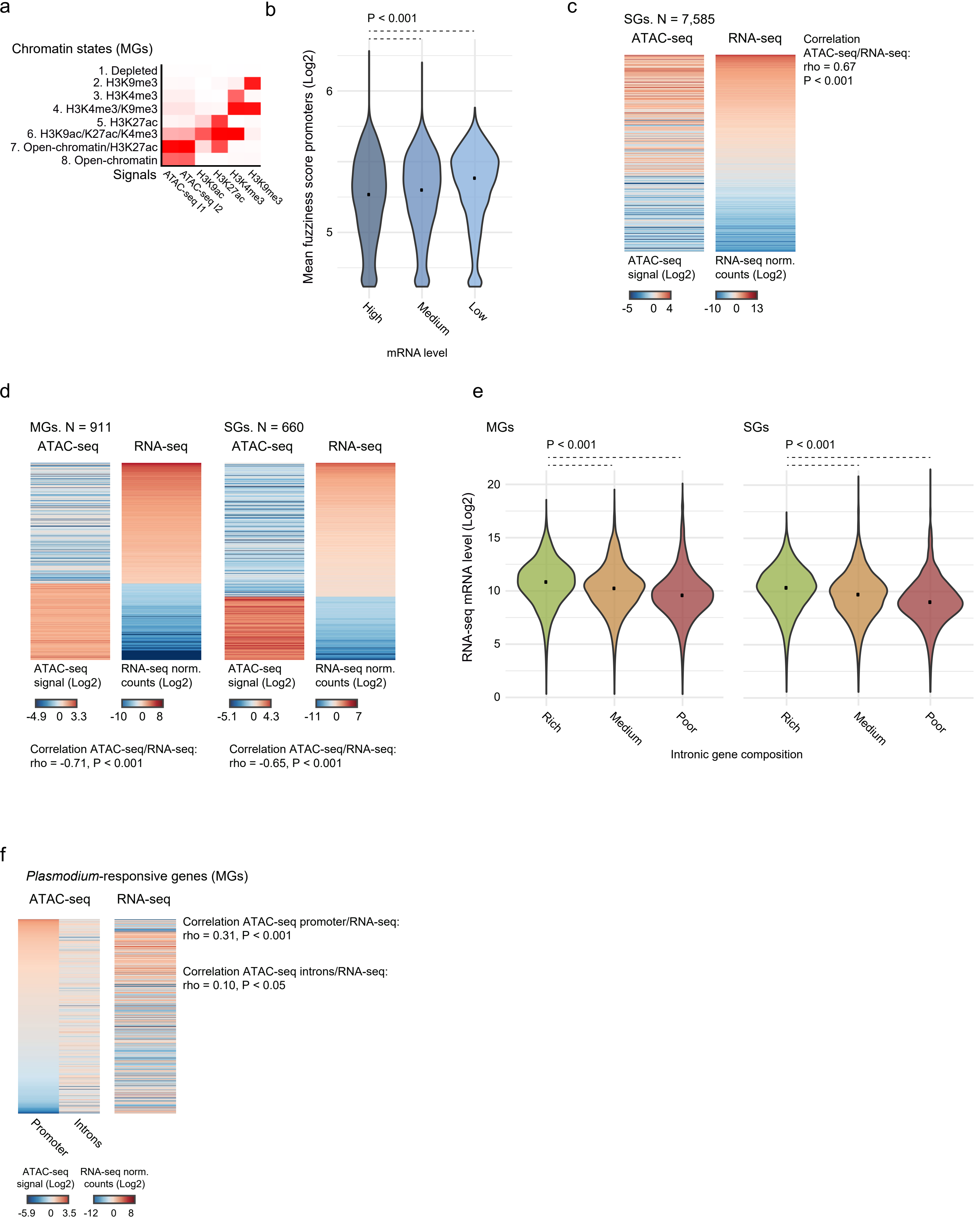


Supplementary Figure 2

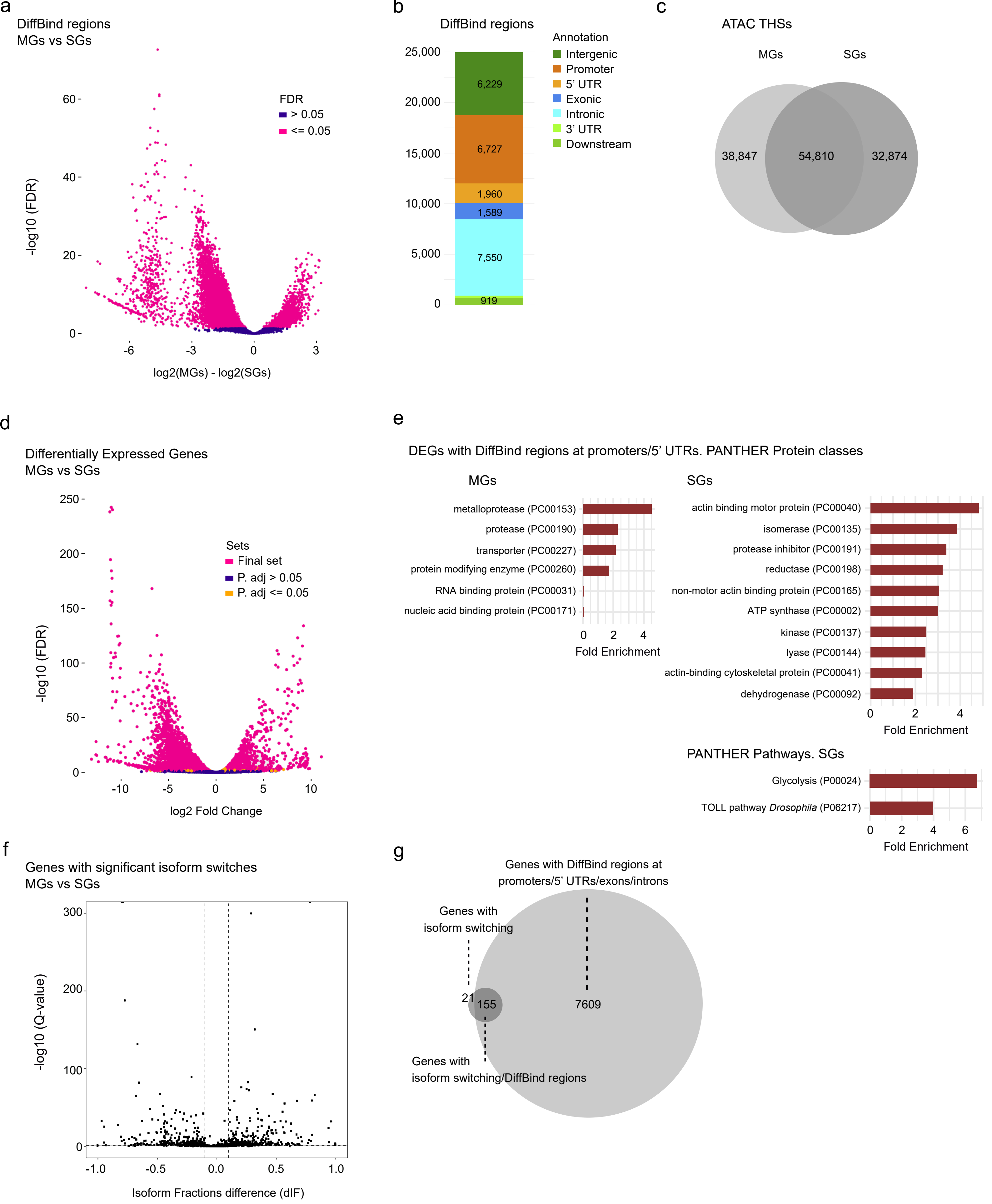


Supplementary Figure 3

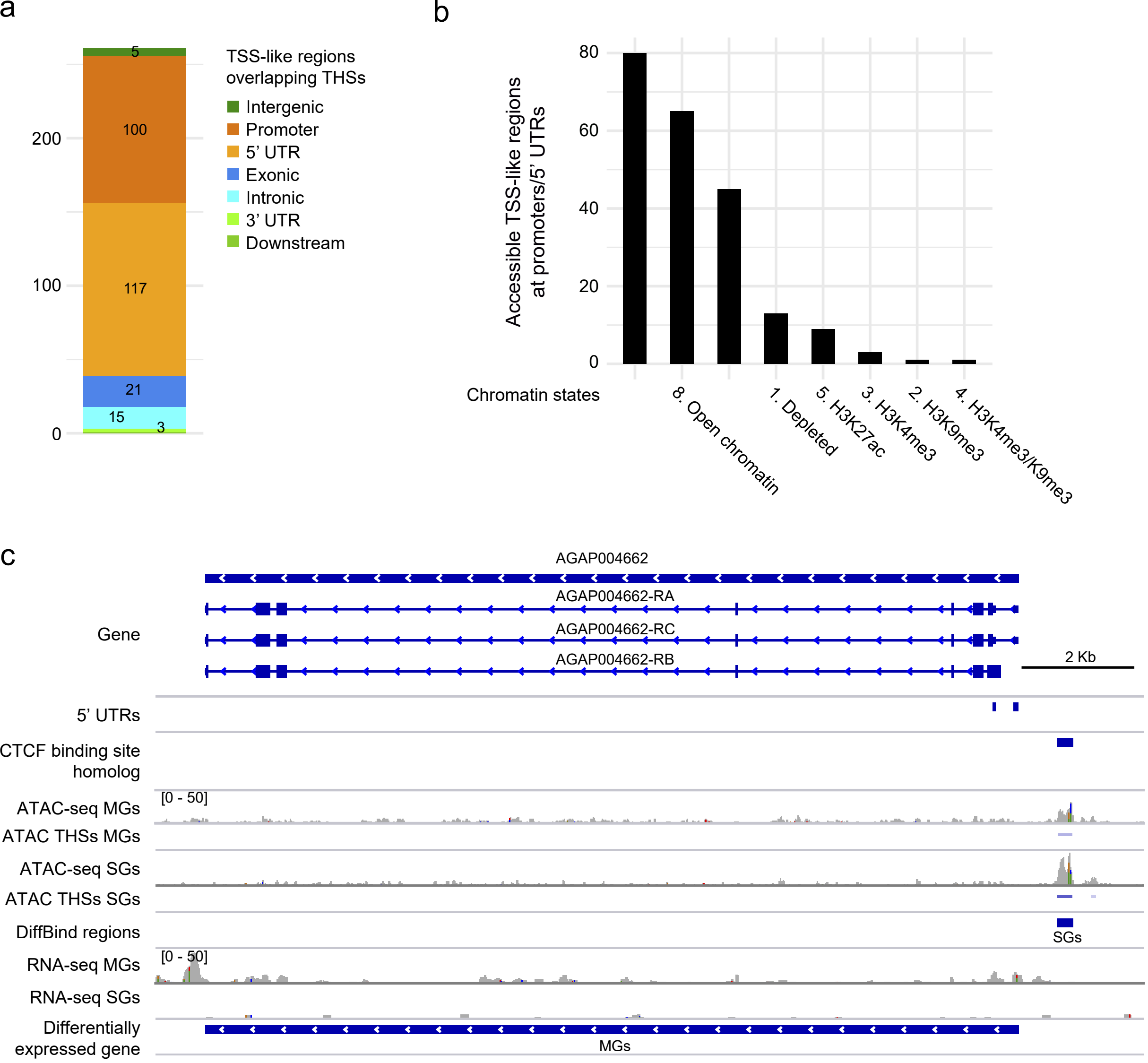


Supplementary Figure 4

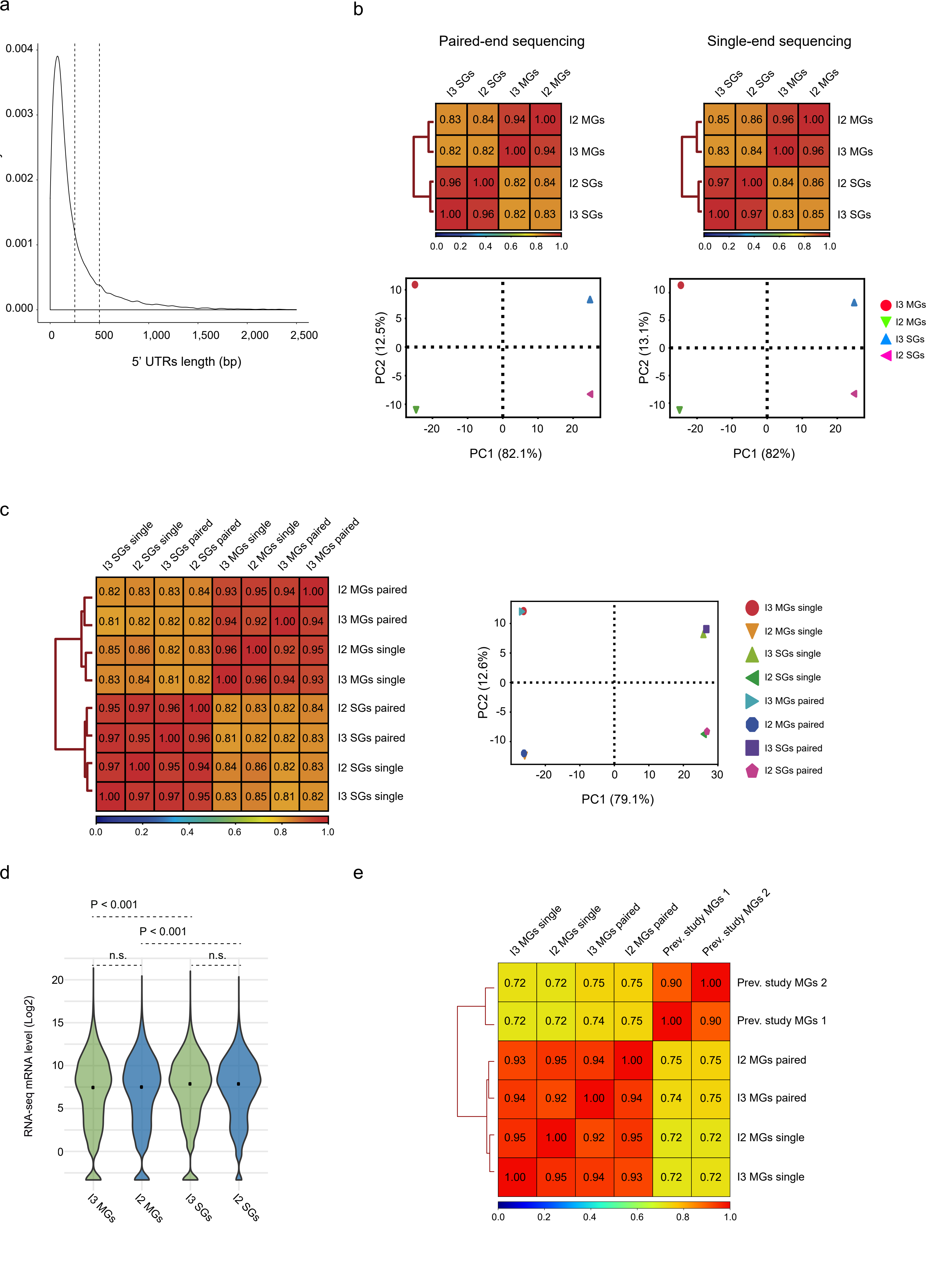


Supplementary Figure 5

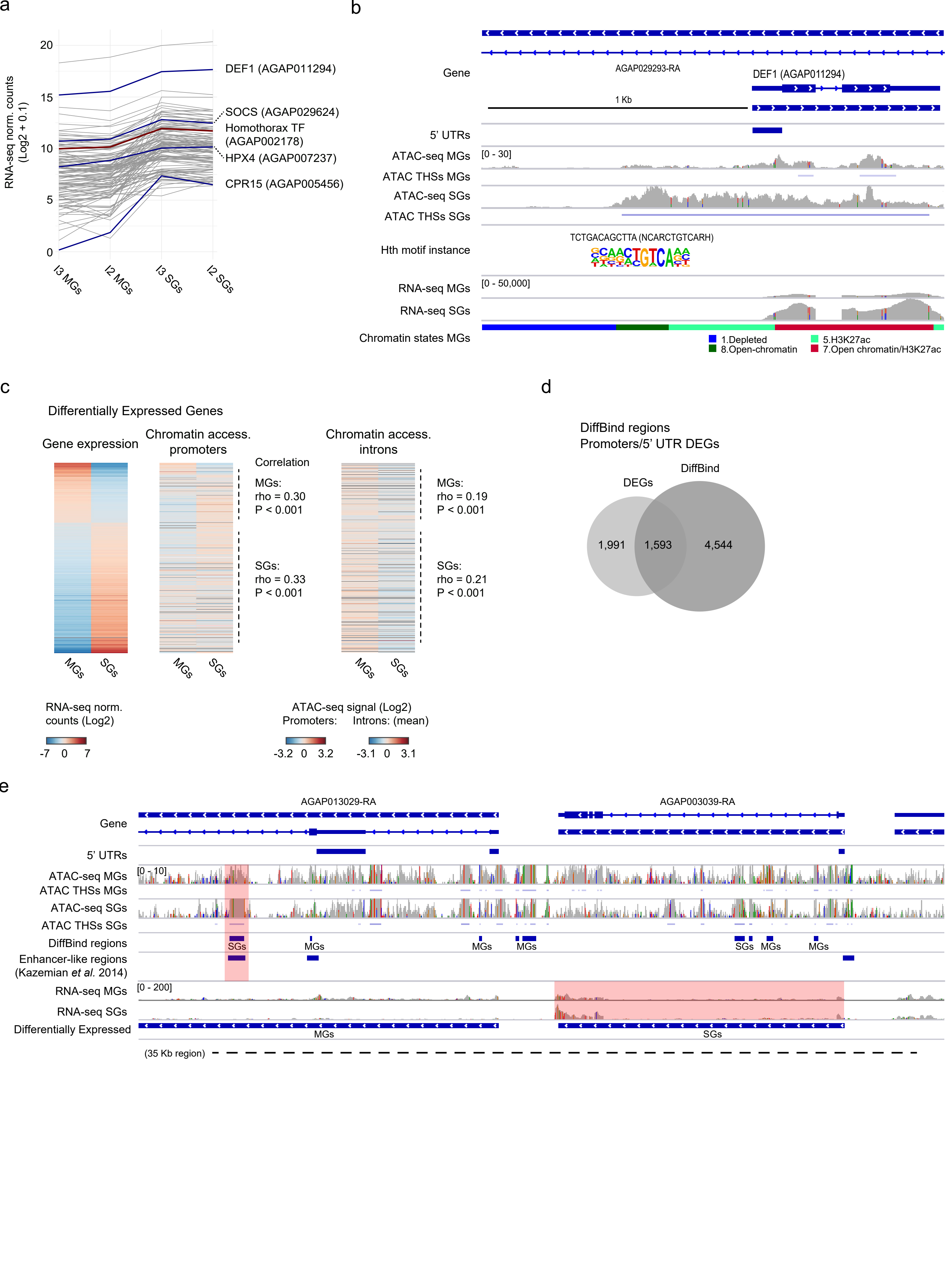


Supplementary Figure 6

### Information on the prevalence and the intensity of infection

| Infections | Prevalence* | Intensity** | Number of mosquitoes | Range of oocyst numbers |
|---|---|---|---|---|
| Infection 1 | 54.5 | 1 | 22 | 0-7 |
| Infection 2 | 75 | 3 | 16 | 0-33 |
| Infection 3 | 57.1 | 1 | 21 | 0-29 |

*Prevalence is the % of infected mosquitoes

**Intensity of infection refers to the median value of oocysts in infected mosquitoes

Prevalence and intensity of infection were both measured at day 7 post-blood meal

### Supplementary Figures

### Supplementary Figure 1

(a) Spearman correlation plot showing similarity between replicates (I1/I2) and tissues: midguts (MGs) and salivary glands (SGs). Samples are ordered by an unsupervised hierarchical clustering approach and cluster by tissue rather than biological replicate (experimental infection). (b) Principal Component Analysis on the replicates (I1/I2) and tissues, MGs and SGs. The plot shows the percentage of variance explained by PC1 and PC2 axes and the distribution of the samples along the first two Principal Components. Note that PC1 explains 96.3% of the variance and PC2 6.2%. Samples cluster by tissue rather than by biological replicate. (c) ATAC-seq density of insertions surrounding *An. gambiae* annotated TSSs (± 1 Kb). The average profiles are shown in solid colored-lines to account for dispersion. Different colors correspond to each biological replicate. Most insertions occur at the TSSs consistently in the two infections. (d) Normalized (RPKM) ATAC-seq nucleosome-free signal at THSs per infection and tissue. Median values are marked with black dots. Chromatin accessibility at THSs is comparable across samples. (e) Venn diagram showing the overlap between THSs identified in each infection for each tissue independently. The areas are proportional to the numbers of regions. The replicability is high as the majority of THSs are present in both experimental infections. (f) Enrichment plots depicting the fraction of ATAC-seq monucleosomal reads (top) and nucleosome-free reads (bottom) in each tissue that map to different genomic features: 5' UTRs, promoters, exons and introns. A larger proportion of ATAC-seq signal occurs at introns and exons. (g) Average profile plot of normalized (RPKM) ATAC-seq nucleosome-free reads at ChIP-seq peaks for various histone post-translational modifications (hPTMs). ChIP-seq peaks are from our previous study and correspond to *P. falciparum*-infected *An. gambiae* midguts[17]. The region plotted comprises ± 1 Kb around the hPTMs peak summits. Profiles in grey represent enrichment at random genomic coordinates. ChIP-seq peaks of H3K9ac, H3K27ac, and to a less extent H3K4me3 are more enriched in nucleosome-free reads (i.e. accessible), whereas H3K9me3 peaks appear depleted.

**Supplementary Figure 2**

(a) Heatmap of ChromHMM emission parameters using an eight chromatin states model based on ATAC-seq and hPTMs enrichment patterns in midguts. The integration of ATAC-seq and ChIP-seq data allows for the partition of the genome into eight epigenetic states. ChIP-seq data was obtained from our previous study[17]. Darker red indicates higher enrichment of a particular signal. (b) Average nucleosome fuzziness score at promoters of genes grouped by gene expression levels (high, medium, low). Nucleosome fuzziness by DANPOS2 measures the deviation of nucleosome positions, and in our study, it is higher for less expressed and silent genes. Plot width accounts for the density of repeated values in the range. Median values are marked with black dots. The P value is indicated for the Wilcoxon signed-rank tests between scores of genes expressed at different levels. (c) Heatmap showing ATAC-seq nucleosome-free enrichment at promoters and gene expression levels of the annotated gene. Data correspond to a subset of non-overlapping genes with a THSs annotated and in which there is a positive relationship between accessibility and gene expression, i.e. transcriptional activation (see Methods). The plot is for salivary glands. Genes are ordered by mRNA levels. Data are mean-centered. ATAC-seq enrichment at promoters is normalized (RPKM). (d) Heatmap showing ATAC-seq nucleosome-free enrichment at promoters and gene expression levels of the annotated gene. Data correspond to a subset of non-overlapping genes with a THSs annotated and in which there is an opposite relationship between accessibility and gene expression, possibly pointing to repressor regulatory events (see Methods). Genes are ordered by mRNA levels. Data are mean-centered. ATAC-seq enrichment at promoters is normalized (RPKM). (e) The plot shows mRNA levels for genes grouped by their intronic composition (see Figure 2d). Intron-rich genes tend to be more expressed. Plot width accounts for the density of repeated values in the range. Median values are marked with a black dot. The P value is indicated for the Wilcoxon signed-rank tests between genes expressed at different levels. (f) Heatmap showing ATAC-seq nucleosome-free enrichment at promoters and introns and gene expression levels of *Plasmodium*-responsive genes[17]. The plot is for midguts. Genes are ordered by ATAC-seq enrichment at promoters. ATAC-seq enrichment is normalized (RPKM). Data are mean-centered.

**Supplementary Figure 3**

(a) Volcano plot displaying DiffBind regions between midguts and salivary glands. The x axis is the fold change difference between tissues, and the y axis indicates the level of significance. Significant DiffBind regions are marked in pink. A negative fold change (x axis) indicates higher accessibility in salivary glands whereas a positive value indicates the opposite, that accessibility is higher in midguts. (b) Annotation of DiffBind regions to features genome-wide: intergenic regions, promoters, 5’ UTRs, exons, introns, 3’ UTRs and downstream regions. Most DiffBind regions annotate to introns. (c) Venn diagram showing the overlap between THSs identified in midguts and salivary glands. The majority of THSs are common, but there is a fraction (30.7% for midguts and 26% for salivary glands) that is tissue-specific. The areas of the circles are proportional to the numbers of THSs. (d) Volcano plot displaying differentially expressed genes (DEGs) between midguts and salivary glands. The x axis is the fold change difference between tissues, and the y axis indicates the level of significance by DESeq2. Significant DEGs are marked in yellow (DESeq2) and in pink if part of the final set (DESeq2/edgeR/DREAMSeq, see Methods). A negative fold change (x axis) indicates higher accessibility in salivary glands whereas a positive value indicates the opposite, accessibility is higher in midguts. (e) Top PANTHER protein classes overrepresented in the set of DEGs with DiffBind regions located at promoters, 5' UTRs and/or introns per tissue. Overrepresented pathways are also shown for salivary glands (results are not significant in the case of midguts). (f) Volcano plot displaying genes with significant isoform switches between midguts and salivary glands. The y axis is the significance while the x axis represents the Isoform Fractions difference (dIF). Dashed vertical lines mark a threshold in significance of 0.05 (y axis) and in dIF of 0.1 in each sense that we use to filter high confidence switching events. A negative dIF indicates higher expression of the isoform in salivary glands, whereas a positive value indicates the opposite, higher isoform mRNA levels in midguts. (g) Venn diagram showing the overlap between genes displaying isoforms switching and genes with a DiffBind region annotated. The areas of the circles are proportional to the numbers of genes.

**Supplementary Figure 4**

(a) Annotation of accessible TSS-like regions to various genomic features: intergenic regions, promoters, 5' UTRs, exons, introns, 3' UTRs and downstream regions. The majority of regulatory sites are located at 5' UTRs and promoters. (b) Chromatin states at the accessible TSS-like regions. As expected, these regulatory regions are accessible and H3K9ac/H3K27ac/H3K4me3-enriched. (c) Chromatin accessibility and gene expression profiles in the region containing the gene encoding the homeobox protein abdominal-A homolog (AGAP004662). Here, this gene is differentially expressed and displays a differentially accessible region between tissues, which is coinciding with a region that is homolog to a CTCF binding site in *Drosophila* at the promoter of the ortholog gene. The tracks shown are for midguts and salivary glands from infection 2. All tracks are shown at equal scale. The location of various genomic features: 5’ UTRs, THSs, DiffBind regions, and the regions.

**Supplementary Figure 5**

(a) Density plot showing the lengths of the annotated 5’ UTRs. The mean length is 253 bp and the higher density occurs around 100 bp, so we expect a 1 Kb window to capture well the promoter regions. (b) Spearman correlation plots (top) and Principal Component Analyses (bottom) for two experimental infections and for paired-end (left) and single-end (right) sequencing approaches. Samples are ordered by an unsupervised hierarchical clustering in the correlation heatmaps. Samples cluster by tissue rather than experimental infections. (c) Plots are as shown in (b) but with the samples by both sequencing approaches. Samples cluster by tissues and experimental infections rather than sequencing approaches. (d) Gene expression levels at each sample. Median values are marked with a black dot. Gene expression is comparable between experimental infections and higher in salivary glands. The P value is indicated for the Wilcoxon signed-rank tests. (e) Spearman correlation plot as shown in (b,c) but integrating the RNA-seq data sets of *P. falciparum*-infected *An. gambiae* midguts in our previous study[17]. The RNA-seq assays are comparable, with around 70% positive correlation.

**Supplementary Figure 6**

(a) Example of Gene Regulatory Network showing groups of co-regulated genes, genes that display active binding sites for a particular TF, and the TF-encoding gene. The line plot shows in dark red the gene expression profile of the homothorax TF-encoding gene (AGAP002178) and the predicted target genes following the same pattern of expression in dark grey. Relevant target genes involved in mosquito immunity are depicted in dark blue. (b) Chromatin accessibility and gene expression profiles in the region containing the DEF1-encoding gene (AGAO011294), which is a *Plasmodium*-responsive gene based on our previous study[17] and a case example here of a gene involved in mosquito immunity that is differentially accessible and expressed between tissues. The tracks displayed are for midguts and salivary glands from infection 2. The location of 5' UTRs, THSs, and the chromatin states are indicated by colored bars. We also show the predicted binding site and motif for the homothorax TF based on the motif enrichment analysis (see Supplementary Figure 6a). All tracks are shown at equal scale. (c) Heatmap showing gene expression levels and ATAC-seq nucleosome-free enrichment at promoters of differentially expressed genes (only for infection 2). There is a positive and significant correlation between chromatin accessibility and gene expression for high/medium expressed genes. Genes are ordered by mRNA levels. ATAC-seq enrichment is normalized (RPKM). Data are mean-centered. (d) Venn diagram showing the overlap between the set of differentially expressed genes and genes with differentially accessible regulatory (DiffBind) regions located at 5' UTRs or promoters. The areas of the circles are proportional to the numbers of genes. (e) Chromatin accessibility and gene expression profiles in the region containing the cation transporter-encoding gene AGAP003039, which is an example of a gene targeted by a distal enhancer[2]. The enhancer region is differentially accessible and more accessible in salivary glands and the target gene is differentially expressed in the same tissue. The tracks shown are for midguts and salivary glands from infection 2. All tracks are shown at equal scale. The location of 5' UTRs, THSs and DiffBind regions are indicated by colored bars. The accessibility at the enhancer regions and the expression profile of the target gene are highlighted in red.

### Supplementary Tables

#### Supplementary Table 1_Infections

Information on the prevalence (percentage of infected mosquitoes), the intensity (mean number of oocysts), the numbers of mosquitoes and the range of oocysts. Samples from Infection 1 (I1) and Infection 2 (I2) are used in the ATAC-seq assay and samples from Infection 2 and Infection 3 (I3) are used in the RNA-seq assay.

#### Supplementary Table 2_Reads alignment statistics

Summary of ATAC-seq and RNA-seq alignment and quality control statistics.

#### Supplementary Table 3_Custom primers

Custom primer sequences specific to ribosomal sequences used in the ribodepletion step.

#### Supplementary Table 4_ATAC-seq THSs

a. Summary of the THSs. b. List of THSs for each tissue and experimental infection annotated to genes and genomic features. Columns include data on the MACS2 peak-calling, whether the THSs are present in both infections, and the chromatin state, overlapping ChIP-seq peaks and whether the annotated genes are *Plasmodium*-responsive in our previous work[17]. Table is ordered by MACS2 Fold Change.

#### Supplementary Table 5_ATAC-seq promoters RNA-seq genes

a. Normalized (RPKM) ATAC-seq enrichment at promoters and gene expression levels of the corresponding gene for each tissue and experimental infection. Columns include gene expression levels by RNA-seq, ChIP-seq enrichment[17], mean nucleosome fuzziness by DANPOS2 and whether the genes are *Plasmodium*-responsive in our previous work[17]. b. Gene structure data and categorization based on the intronic/exonic composition.

#### Supplementary Table 6_Differential analyses

a. Differential chromatin accessibility regions (DiffBind) between tissues. Columns include data on the DiffBind analysis, the annotation to genomic features, the annotated THSs, gene expression levels by RNA-seq, and coincidence with ChIP-seq peaks and chromatin states in our previous study[17]. Table is ordered by DiffBind Fold binding affinity. b. Differentially Expressed Genes (DEGs) between tissues. Columns include data by DESeq2, but the genes are the result of combining DESeq2/edgeR/DREAMSeq. c. Integration of DiffBind regions (a) annotated to DEGs (b). d. Results of the GO terms, protein classes and pathways PANTHER overrepresentation tests (see Results).

**Supplementary Table 7_ATAC-seq nucleosome dyads**

a. Summary of the nucleosome dyads. b. List of predicted nucleosome dyads for each tissue and experimental infection. Occupancy levels by NucleoATAC and coincidence with the ChIP-seq peaks in our previous study[17] are included. Table is ordered by NucleoATAC occupancy.

**Supplementary Table 8_Chromatin states segmentation**

Segmentation of the genome in chromatin states by ChromHMM based on midguts ATAC-seq and ChIP-seq[17] data.

**Supplementary Table 9_Isoform switches analyses**

Genes switching isoforms expression between tissues with annotated DiffBind regions.

**Supplementary Table 10_DiffBind regions motifs**

List of enriched consensus motifs corresponding to know *Drosophila* TFs in the set of THSs at DiffBind regions that annotate to the promoters or 5' UTRs, or introns, of differentially expressed genes. Results are separated by tissues and activator and repressor candidates.

**Supplementary Table 11_Novel regulatory regions**

a,b. *Anopheles* enhancers previously predicted by others[4] (a) and novel enhancer-like regions predicted here based on *D. melanogaster* homology[48] (b). The overlapping THSs and annotated genes and whether the enhancers were experimentally validated in previous studies are included. c. TSS-like regions predicted here based on *D. melanogaster* homology. The overlapping THSs, annotated genes and whether the genes have previously annotated 5’ UTRs are included. d) *An. gambiae* regions homolog to *Drosophila* CTCF binding sites[49]. The overlapping THSs and annotation are included.

**Supplementary Table 12_Enhancer region motifs**

List of enriched consensus motifs corresponding to known *Drosophila* TFs in the set of THSs at *An. gambiae* enhancer-like regions.

**Supplementary Table 13_Motifs activating repressor**

List of enriched consensus motifs corresponding to know *Drosophila* TFs in the set of THSs that annotate to the promoters of active genes, and low expressed or silent genes.

**Supplementary Table 14_Motif instances hth, srp, Trl**

Enriched motif instances at different sets of THSs (see Supplementary Results) for the TFs: homothorax (a), serpent (b) and trithorax-like (c).

### Supplementary Results

#### Motif analyses of accessible regions in different tissues

For the set of THSs that annotated to the promoters of actively transcribed genes (4,197 for MGs, and 5,059 for SGs; Supplementary Table 4), we conducted DNA-binding motif enrichment analyses to find enriched *de novo* motifs matching consensus TFs sequences (binding sites). Supplementary Table 13 shows the list of novel motifs and the similarity to known *Drosophila* TF binding sites. Among the top hits, we predicted the vismay, homothorax, or trithorax-like TFs, which are known to be involved in processes related to development, morphogenesis and signaling (Supplementary Table 13). For motifs found in THSs-annotated promoters, we identified clusters of co-expressed genes that appeared to be regulated by the same set of TFs, i.e. Gene Regulatory Networks (GRNs). For example, this is the case of the *An. gambiae* gene AGAP002178, which is ortholog to the *Drosophila* homeobox TF homothorax. We found binding motifs for this predicted TF at the promoters of over 100 genes, including immune-related genes, such as the defensin antimicrobial peptide *def1* (Supplementary Figure 6a and Supplementary Table 14). Both the target genes and the TF-encoding gene shared similar transcriptional profiles (Supplementary Figure 6b and Supplementary Table 5). In addition to regulatory regions annotated to active genes, a small portion of THSs annotated to weakly-expressed and silent genes, and thus may correspond to TF binding sites with potential repressor roles (MGs: 389 THSs, SGs: 285 THSs). In this case, the motif analyses predicted *Drosophila* TFs with known repressor roles, including buttonhead, caudal or hunchback (Supplementary Table 13). By restricting the analysis to THSs that annotate to the promoters of immune-related genes[1], we also predicted *Drosophila* TFs with relevant roles in immune response such as dorsal, dif, or Deaf1 (Supplementary Table 13).

**Tissue-specific chromatin accessibility correlates with differential gene expression**

To investigate the functional consequences of differential regulatory events, we first performed gene expression analyses that result in a set of 3,584 Differentially Expressed Genes (DEGs): 68.6% more expressed in SGs and 31.4% in MGs (Supplementary Figure 3d and Supplementary Table 6). The integration of the ATAC-seq and RNA-seq data for genes that appeared differentially expressed between the two tissues (3,584 DEGs; Supplementary Figure 3d and Supplementary Table 6), revealed a significant correlation between the level of accessibility at the regulatory sites (promoters or introns) and the levels of gene expression in the same tissue (Spearman test, promoters: rho 0.30, P < 0.001 (MGs); rho 0.33, P < 0.001 (SGs); introns: rho 0.19, P < 0.001 (MGs); rho 0.21, P < 0.001 (SGs); Supplementary Figure 6c).

Around 33% of the DiffBind regions (8,197) annotated to 70% of the DEGs (2,419), and 27.6% of these (2,259) were located at the 5' UTRs or the promoters of 1,593 DEGs (Supplementary Figure 6d). For around 70% of the DEG/DiffBind overlapping regions, we also observed that the change in the ATAC-seq and gene expression patterns between tissues happened in the same direction. That is, it seemed higher accessibility tends to be linked to higher gene expression in the corresponding tissue (Supplementary Table 6).

**Chromatin accessibility allows for the identification of novel *An. gambiae* cis-regulatory elements**

In *An. gambiae*, previous studies have computationally predicted a total of 1,679 enhancers based on a machine learning approach[2], by searching for TF motifs in *Drosophila* that are bound by enhancers[3] or by STARR-seq[4]. Of these, only 5 have been experimentally validated in *An. gambiae*[2], and 6 in *An. coluzzi*[4]. When crossing the 1,679 *An. gambiae* and 6 *An. coluzzi* enhancers with our ATAC-seq data set, we found that 42% (708/1,685) overlapped with THSs (1,889), and thus may correspond to active enhancers (Supplementary Table 11). In agreement, this list of enhancer candidates captured 4 out of the 5 previously experimentally validated *An. gambiae* enhancers.

Compared to the number of publicly available enhancers for model organisms, such as *D. melanogaster,* the number of predicted enhancers for *A. gambiae* is relatively low. Enhancers are sequences that may be subjected to rapid evolution due to their functional implications[5, 6, 7, 8], but it is anticipated to find some degree of sequence conservation between phylogenetically close metazoans, including dipterans[9, 10, 11, 12]. Thus, we wondered whether we could use public updated sets of *Drosophila* enhancers in combination to our chromatin accessibility map to further expand our set of mosquito cis-regulatory elements identified by ATAC-seq. To this end, we used the recently updated EnhancerAtlas v2.0[13, 14] to download more than 40,000 *Drosophila* enhancer elements for which a target gene has been predicted. In addition, we added to our working set 783 *Drosophila* enhancer-target gene pairs that have been previously characterized and experimentally validated[15]. Taking advantage of the high homology between *D. melanogaster* and *An. gambiae*, we used the LiftOver software[16] to create a map of candidate enhancers in mosquitoes. This approach resulted in 1,126 *An. gambiae* enhancer-like regions, most of which (1,122) were potentially new, that is, they do not coincide with previously predicted mosquito enhancers. By intersecting the ATAC-seq data, we observed that around 10% (103) of these putative enhancer-like regions overlapped with 383 THS regions (Supplementary Table 11), and 27 of these have been experimentally validated in *Drosophila*[15].

Next, putting the two sets of accessible regulatory sequences by ATAC-seq together we built a final set of 811 *An. gambiae* enhancer-like regions, which overlapped with 2,272 THSs and annotated to 563 genes (Supplementary Table 11). For around 80% (633) of the accessible enhancer-like regions by ATAC-seq that we annotated to the closest gene, that gene coincided with the target reported previously by others[2, 4, 15]. This would indicate these are proximal enhancer elements. In contrast, 20%

(167) of the enhancer sites did not annotate to the closest gene based on target homology analysis, which would point to distal regulation. However, of these, we report 115 in which the original target predicted by others (either in *D. melanogaster* or *An. gambiae*) annotates to multiple *An. gambiae* orthologs (Supplementary Table 11). We also report 11 enhancer-like sequences that were originally annotated to target genes in *D. melanogaster* without an *An. gambiae* ortholog. As a result of these filters, we ended up with 52 that could be considered as potential distal enhancers. Out of the 39 that were originally transferred from computationally predicted *An. gambiae* enhancers[2], 31 could be considered distal in this study (Supplementary Table 11). This is the case of the cation transporter gene AGAP003039 (Supplementary Figure 6e). Similarly, out of the 13 potentially distal enhancers transferred from *D. melanogaster*[15], we observed 9 that were also distal in *Drosophila*: 3 appeared located in the same chromosome than the target genes, and 6 were located in different chromosomes. Contrary, there are 4 that were reported as proximal in *Drosophila* but we observed that in the mosquito the enhancer sequence and the ortholog target genes are located in different chromosomes.

Among the *An. gambiae* genes harboring accessible enhancer-like regions, we found 13 immune-related genes[1] (Supplementary Table 11). These included the cytokine *spz3* (AGAP008360) or the C-type lectin *ctlga1* (AGAP010196). Others were components of the cytochrome P450, such as the *cyp49a1* or the *cyp6y2*, or TFs, such as the remodeling and spacing factor 1 (AGAP001386) and the enhancer binding protein AGAP011096 (Supplementary Table 11). We also identified enhancer candidates for 76 *Plasmodium*-responsive genes[17], such as the cytochrome P450 component *cyp6m4* (AGAP008214), the enhancer of polycomb protein (AGAP008026) or the CCAAT/enhancer binding protein (C/EBP) (AGAP011096), as well as genes encoding for various histone acetyl- and methyl-transferases (Supplementary Table 10).

Regarding the integration of the differential gene expression data and differential chromatin accessibility at enhancer-like regions, one third (33%) of the proximal enhancer-like regions (211 out of 633) coincided with DiffBind regions. Of these, 51 were annotated to DEGs and we showed the most frequent role was activating: in 37 cases the regulatory region was more accessible in the same tissue where the gene was more expressed, and in 14 cases the relationship was the opposite, which points to a repressor function.

The current *An. gambiae* annotation includes 5’ UTRs and TSSs for the majority of genes, but around 20% of them (2,890 out of 13,057) do not have annotated 5’ UTRs. In this study, we observed a higher density of THSs at 5’ UTRs or ATGs (see above), so we used our chromatin accessibility map to find novel *An. gambiae* TSS-like elements. TSSs tend to be highly conserved and collections of *D. melanogaster* TSSs are also publicly available, so same than above, to propose new mosquito TSS-like elements we used a dataset of 16,792 *Drosophila* TSSs from the Eukaryotic Promoter Database[18]. Following the same homology approach, we identified 917 TSS-like sites in *An. gambiae* that annotated to 819 genes (Supplementary Table 11). For 34.5% (316/917), the annotated gene was a direct ortholog to the original *Drosophila* gene (Supplementary Table 11).

### Supplementary Methods

#### External data

We used ChIP-seq data sets for various post-translational modifications of histones (hPTMs) in *P. falciparum*-infected *An. gambiae* midguts (MGs) obtained from our previous study[17]. To compute enrichment we calculated normalized counts using

BEDTools *intersect* -c[19] (v2.29.2). The signal was input-corrected (ratio), RPKM-normalized and a pseudocount added when needed to get finite values. We tested the degree of replicability in the gene expression estimates, by measuring the correlation between RNA-seq data from this study and our previous study. Based on the high correlation we used the list of *Plasmodium*-responsive genes in the ATAC-seq analysis.

**Libraries and mapping QC**

Alignment statistics are included in Supplementary Table 2. We performed quality control on raw reads by FastQC (http://www.bioinformatics.bbsrc.ac.uk/projects/fastqc, v0.11.9). More statistics were obtained using QualiMap[20] (v2.2.2d). Once the ATAC-seq reads were mapped and filtered, and following the ENCODE guidelines, we measured library complexity using the Non-Redundant Fraction (NFR) and the PCR Bottlenecking Coefficients (PBC1, PBC2). After ATAC-seq peak calling (see next section), we further checked quality with the Fraction of Reads in Peaks (FRiP) score. We used ataqv[21] (v1.1.1) to plot the fragment length distributions of ATAC-seq reads. Correlation and Principal Components plots, signals and coverage profiles were obtained using deepTools[22](v3.4.1), ngs.plot[23] (v2.6.3) and V-plots (adapted code from https://github.com/jinxu9/ATACseq/blob/master/libs/pyMakeVplot.py).

**RNA isolation, RNA-seq library preparation and sequencing**

We designed custom primers specific to mosquito ribosomal sequences to reduce the percentage of ribosomal sequencing reads in the samples. We performed multiple sequences alignments using Clustal Omega[24] and known rRNA sequences from the updated *An. gambiae* genome to extract representative sequences, which were then used by Nugen Technologies to generate a custom design with unique probes. The list of custom primers is in Supplementary Table 3.

**ATAC-seq data processing and analyses**

We performed ATAC-seq data analyses according to the recommendations by ENCODE Pipeline prototype (https://www.encodeproject.org/atac-seq/), with steps including read trimming, mapping, peak calling and signal generation. For the annotation of the THSs to genomic features, we combined the HOMER[25] (v4.11) and ChIPseeker[26] (v1.22) software. We loaded the AgamP4 reference genome v.2.00 and the AgamP4.12 gene set in HOMER using the *loadGenome.pl* module. THSs annotation was conducted using the *annotatePeaks.pl* module with defaults parameters. The same reference genome was loaded in ChIPseeker using the *makeTxDbFromGFF* R function. The *annotatePeak* R function was run with default arguments, except for the reference genome (TxDb object), the Bioconductor annotation[27] and the coordinates defined as promoter regions (tssRegion = c(-1000,0)). We combined the annotations using both software, prioritizing the overlapping genomic features in the following order: 5' UTRs, 3' UTRs, Promoters, Exonic, Intronic, Downstream, Intergenic.

To predict nucleosome dyads positions we used NucleoATAC[28] (v.0.3.4). First, we obtained a new set of ATAC-seq peaks as described in Methods, but including the --broad flag in peak calling by MACS2. These peaks were then extended 100 bp, merged to get unique coordinates and used as input regions for the NucleoATAC with default parameters following the manual.

Unless otherwise specified, we performed general interval operations including intersect, merge, shuffle or slop were performed using the BEDTools suite. Statistical tests and plots were computed in R using Bioconductor when needed (v3.6). Bar and violin plots were produced using the ggplot2 R package[29] (v3.3).

**RNA-seq data processing and analyses**

For the alignment of the RNA-seq sequencing reads, we first created the genome index using STAR *genomeGenerate* mode with default parameters, except for --genomeSAindexNbases 13, and --sjdbOverhang 150 or --sjdbOverhang 75 for the single-end and paired-end sequencing approaches respectively. Next, we used the AgamP4 reference genome (v.2.00) and the AgamP4.12 gene set available in VectorBase[30] to map the reads with default parameters except for --twopassMode Basic --alignIntronMin 1 --alignIntronMax 249416 --outFilterScoreMinOverLread 0.3 --outFilterMatchNminOverLread 0.3. The reads and alignments statistics are included in Supplementary Table 2.

We obtained RNA-seq counts at the gene level using the CoCo software. CoCo is a counting software designed to quantify expression of nested and multi-mapped genes[31]. First, it creates a corrected annotation controlling for embedded genes. We provided the AgamP4.12 gene annotation to the *correct_annotation* module, manually processed to include gene biotypes, and ran the script with default parameters. Second, to quantify gene expression levels, we used the *correct_count* module with default parameters, except for -s 1 to specify the libraries strandness (strand-specific assay), the -i parameter to specify each library insert size and the -p argument to mark the use of paired-end reads. Raw read counts from both sequencing approaches (single-end and paired-end) were then summed for each sample and were used as input for DESeq2 in order to perform library normalization. The design included the tissue as main factor and the infection as co-variable to control for inter-experiment variability. Genes with low number of counts (<5) were discarded.

**Integration of ATAC-seq, RNA-seq and ChIP-seq data**

We performed genome segmentation in chromatin states using ChromHMM[32] (v.1.2). For the binarization step, we used default parameters except for -b 200 and -paired. We tested ChromHMM with different *a priori* number of states and chose eight as the best number to maximize informative features (i.e. unique combinations of histone modifications and open chromatin and not redundant states). We assigned chromatin states to different regions of interest (i.e. THSs) using BEDTools *intersect*.

Heatmaps showing chromatin accessibility, histone modification enrichment and mRNA levels were built using the iheatmapr R package[33] (v0.4.7). For comparative and visualization purposes, ATAC-seq, RNA-seq and ChIP-seq enrichment data are log2-scaled and mean-centered. Average profile plots representing ATAC-seq and ChIP-seq enrichments (RPKM normalized and input-corrected) centered on TSSs, peaks coordinates or gene bodies were built using ngs.plot. Violin plots, boxplots and barplots were produced using the ggplot2 R package.

**Differential ATAC-seq analyses**

ATAC-seq reads for each infection and tissue were counted using the *dba.count* function with default arguments, except for the score for the normalization step (*DBA_SCORE_TMM_READS_FULL*). The significant differential enrichment test was computed using the *dba.analyze* function that includes the DESeq2 module with default parameters and 0.05 as significance threshold. The THSs provided to DiffBind were automatically merged during the processing and unique regions were analyzed, so each significant differentially accessible region (DiffBind region) corresponded to multiple THSs. We assigned each DiffBind region to the THS that overlapped more than half of it (BEDTools *intersect* -f 0.51) and displayed the highest MACS2 Fold Enrichment score. A high confidence set of DiffBind regions was obtained by filtering out the regions not consistently displaying the same THS in the two infections.

Plots for the visualization of the differential ATAC-seq enrichment analysis were performed using the ggplot2 R package following the DiffBind Bioconductor vignette.

**Differential RNA-seq analyses**

To account for both under-dispersed and not under-dispersed data, the R packages DEseq2/edgeR/DREAMSeq were used independently[34, 35, 36], and the results were combined to obtain a high confidence set of DEGs. In all cases, the matrix of raw reads counts was used as input and the significance threshold was 0.05. DESeq2 and edgeR are both based on negative binomial linear models[34, 35] and their design included condition as the main factor. Genes with lower read counts (<5) were discarded. The DREAMSeq method combines negative binomial and double Poisson models to properly fit a wider range of data[36]. Here, we used default parameters except for specifying the *DREAMSeq.Mix* model and a Fold Change threshold of 1.5 (*fc* argument). The final list of DEGs was obtained by taking the genes present in all three data sets.

**Differential isoform expression analyses**

We used the IsoformSwitchAnalyzeR R package[37] (v1.8) to analyze differential gene isoforms expression between MGs and SGs and to determine underlying alternative splicing events following the Bioconductor vignette. First, the quantification of the expression at the level of gene isoforms was performed using Salmon[38] (v1.1.0). We used *salmon index* with default parameters to create an index using the reference transcript sequences (AgamP4.12, VectorBase). Then we used *salmon quant* with default parameters except for -l A and –validateMappings. The RNA-seq reads provided were the ones after trimming of adapters (BBDuk) and rRNA filtering (SortMeRNA). Next, Salmon count data from both sequencing approaches and the annotation from the AgamP4.12 gene set were imported into IsoformSwitchAnalyzeR. To conduct the analyses, a BSgenome object was produced using the *forgeBSgenomeDataPkg* R function and the AgamP4 v2.00 reference genome. Second, the gene isoforms switching between tissues were determined using the *isoformSwitchAnalysisPart1* wrapper function with default parameters, except for the custom BSgenome object (*genomeObject* argument), a cutoff for finding switches of 0.1 (*dIFcutoff* argument) and the specification of DEXSeq as statistical method (*switchTestMethod* argument). Here, a cutoff of 0.1 means a 10% threshold in isoform usage variation to consider an isoform switching. Sequences for the gene isoforms were also obtained as output (outputSequences = T) to use as input for downstream analyses. The second part of the analysis was the characterization of alternative splicing events underlying gene isoform switching, and it required the manipulation of the transcript sequences. We annotated the sequences obtained above using CPAT[39] to assess their coding potential, Pfam[40] to annotate protein domains, SignalP[41] to annotate signal peptides and NetSurfP-2[42] to analyze the Intrinsically Disordered Regions. We ran these external tools using the available web servers and following the instructions in the Bioconductor package vignette. CPAT was run locally using the default parameters for all the scripts, the AgamP4.12 gene set and the AgamP4 reference genome. The final step was then executed using the *isoformSwitchAnalysisPart2* wrapper function with default parameters except for: dIFcutoff = 0.1, n = NA, removeNoncodinORFs = F, codingCutoff = 0.39, outputPlots = T, consequencesToAnalyze = c('intron_retention', 'coding_potential', 'ORF_seq_similarity', 'NMD_status', 'domains_identified', 'IDR_identified', 'signal_peptide_identified') and pathToCPATresultFile, pathToPFAMresultFile, pathToSignalPresultFile and pathToNetSurfP2resultFile arguments pointing to the results of the external sequence analyses above. The consequences of some isoforms switching were predicted using the function *extractConsequenceSummary* and underlying alternative splicing events were predicted by the *analyzeAlternativeSplicing* function.

**Integration of differential accessibility, differential gene expression and differential gene isoforms switching analyses**

Differentially accessible regions (DiffBind) were annotated to genomic features and to the set of DEGs (DESeq2/edgeR/DREAMSeq). We used Fold Change values as a quantitative measure of the extent and the sense of change of both differential expression and accessibility. For the integration of gene isoforms switching results and differential chromatin accessibility, we isolated the genes with isoform switching with DiffBind regions located at promoters, 5’ UTRs, introns or exons. This allowed the identification of genes showing differences in isoform expression that could be due to differential chromatin accessibility at regulatory regions.

We performed functional analyses and GO term overrepresentation tests of DiffBind-annotated DEGs using the Statistical Overrepresentation test in PANTHER[43] (v14.1, 20190711). Violin plots, boxplots and barplots were produced using the ggplot2

R package. Heatmaps were built using the iheatmapr R package. For comparative and visualization purposes, data were log2-scaled and mean-centered.

**Characterization of novel regulatory elements**

To map active regulatory elements, we first used *An. gambiae* enhancers already predicted by others. In particular, a set of 1,628 enhancers was predicted by Kazemian *et al*[2] using an *in silico* machine learning approach and a *D. melanogaster* training set of cis-regulatory modules involved in developmental patterning. Only 5 of these were validated *in vivo* by transgenic reporter assays. Ahanger *et al*[3] predicted 51 enhancer elements on the Hox gene complex by looking for motifs of two *Drosophila* TFs. Nardini *et al*[4] identified by STARR-seq, and experimentally validated by luciferase assays, 6 enhancers in *An.coluzzi*/*An.gambiae*. Next, to expand the set of mosquito enhancers, we used the UCSC LiftOver tool webserver to transfer enhancer coordinates between *D. melanogaster* and *An. gambiae*. We mapped the enhancers from the *D. melanogaster* dm3 or dm6 genome to the *An. gambiae* anoGam3 genome. The regions at the chromosomes chrM and chrUn were excluded. The anoGam3 genome is not the *An. gambiae* most updated version and it corresponds to the AgamP3 version at VectorBase. To transfer the regions from the anoGam3 to the AgamP4 v2.00 genome available at VectorBase, we obtained the necessary chain files using the flo pipeline[44] (https://github.com/wurmlab/flo) and we provided them as input to the LiftOver command line version with default parameters. We used the ABC pipeline[45] (v0.2) following the manual and our ATAC-seq data and H3K27ac previous data set[17] to compute an activity score based on accessibility and H3K27ac enrichment. We divided the predicted enhancer activity into high, medium, low groups (*Hmisc::cut2* R function).

When needed to compare the annotated genes in the two organisms, we used FlyBase[46] (FB2019_03), VectorBase[30] (Release VB-2019-08) and OrthoMCL[47] to get orthologs between the species.

**Motif enrichment analysis**

For *de novo* motif analysis using HOMER, we used the *findMotifsGenome.pl* module. We limited the number of background sequences to double the number of THSs analyzed using the argument -N. The rest of parameters were by default, except for: -len 8,10,12 -mset insects -preparse. Only motifs enriched in more than approximately 2-3% of the target sequences and below a threshold P value of 10E−10 were considered, and we avoided results corresponding to low complexity motifs and offsets or degenerate versions of highly enriched motifs. We used the *annotatePeaks.pl* module with the argument -mbed to find motif occurrences and instances in each subset of THSs.
